# A Design Principle for slow-wave sleep firing pattern with Na^+^ dynamics

**DOI:** 10.1101/2024.08.13.607682

**Authors:** Tomohide R. Sato, Koji L. Ode, Fukuaki L. Kinoshita, Hiroki R. Ueda

## Abstract

Non-rapid eye movement (NREM) sleep is characterized by electroencephalography (EEG) signals with high amplitude and low frequency. This signal is thought to originate from the synchronized activity of cortical neurons, showing the alternating bursting state (up state) and resting state (down state). This activity is termed as slow-wave sleep (SWS) firing pattern. We previously proposed the importance of Ca^2+^-dependent hyperpolarization pathway in generating this firing pattern by introducing the averaged-neuron (AN) model, which describes neuronal activity based on the Hodgkin-Huxley type model. In the AN model, Ca^2+^-dependent K^+^ channels are involved in the transition from the up to the down state. Here we focus on the intracellular Na^+^ dynamics which are not explicitly described in the AN model. A revised AN model, termed as Na^+^-centered AN (NAN) model, proposes that the activation of voltage-gated Na^+^ channels leads to intracellular Na^+^ accumulation, which in turn triggers the activation of Na^+^-dependent K^+^ (KNa) channels or Na^+^/K^+^ ATPases, resulting in the down state. Changes in the activation kinetics of voltage-gated Na^+^ channels are important in shaping SWS firing pattern as well as explaining the inter-spike interval changes between SWS and AWAKE firing pattern. Mathematically, transition from the up state to the down state occurs in accordance with the change in the number of the fixed point in the dynamical system with the changes in the intracellular Na^+^ concentration. The importance of Na^+^-dependent pathway is elucidated even with the coexistence of Ca^2+^-dependent pathway. Subsequent analysis with network model suggests that the result of averaged neuron model with Na^+^ pathway can be extended to the population of neurons. Therefore, our model proposes that voltage-gated Na^+^ channels and Na^+^-dependent K^+^ channels or Na^+^/K^+^ ATPases are also the candidate pathways for the generation of SWS firing pattern.

## INTRODUCTION

Electroencephalography (EEG) is used to observe ensemble activity of cortical neurons during sleep and awake states. Previous studies showed that a high-amplitude and low frequency EEG pattern is observed during non-rapid eye movement (NREM) sleep^1^. This EEG activity is represented as slow-wave activity. When slow-wave activity is perceived, a characteristic electrophysiological firing pattern is detected in cortical neurons, which is composed of two states; a depolarized bursting phase (up state) and a hyperpolarized resting phase (down state)^2,3^. Hereafter, we call this pattern slow-wave sleep (SWS) firing pattern.

Several computational models succeeded in recapitulating the SWS firing pattern^4-6^, yet molecular details of this firing patterns are still elusive. A study succeeded in generating SWS firing pattern by using the averaged neuron (AN) model^7^. The AN model describes neuronal firing based on the Hodgkin-Huxley model^8^ and incorporated 13 components including receptors of neurotransmitters, ion channels and Ca^2+^ pumps. The AN model leveraged the concept of mean-field approximation and enabled detailed analyses of individual components. Later study constructed a simplified version of AN model (SAN model)^9^ to conduct a mathematical analysis about the mechanisms underlying the SWS firing pattern. In the AN/SAN models, Ca^2+^-dependent hyperpolarizing currents mediated by Ca^2+^-dependent K^+^ (KCa) channels mainly contribute to induce the down state. On the other hand, the role of Na^+^-dependent K^+^ (KNa) channels is indicated in the previous study^10^. KNa channels are supposed to regulate the resting membrane potential and the adaptation of firing rate^11,12^. The activation of KNa channels is dependent on the intracellular Na^+^ concentration. The increase of the intracellular Na^+^ concentration is typically mediated by voltage-gated Na^+^ channels. However, whether activation of voltage-gated Na^+^ channels serves to induce hyperpolarizing down state in SWS firing pattern remains elusive: this is because voltage-gated Na^+^ channels which activate KNa channels also create depolarizing currents.

Typical voltage-gated Na^+^ channels are completely inactivated within a few milliseconds, but residual Na^+^ currents are observed, named the persistent Na^+^ currents^13^. Electrophysiological measurement showed that the persistent Na^+^ currents are observed because Na^+^ channels are activated at a lower voltage and inactivation of the channels are much slower than Na^+^ channels described in Hodgkin-Huxley model^14^.

On the other hand, a major factor responsible for Na^+^ efflux is Na^+^/K^+^ ATPases. Theses ATPases exchanges intracellular three Na^+^ and extracellular K^+^ by consuming the energy of ATP hydrolysis^15^. This activity is enhanced when intracellular Na^+^ is elevated^16^. Na^+^/K^+^ ATPases usually work to restore the concentration gradient of Na^+^ and K^+^ between inside and outside of the cells, which is counteracted by passive diffusion during the occurrence of the action potential.

Na^+^ dependent hyperpolarization pathway and Ca^2+^ dependent hyperpolarization pathway can similarly work in the transition from the up state to the down state. However, such Na^+^- or Ca^2+^-dependencies with different influx current, efflux current, and baseline concentrations of intracellular Na^+^ and Ca^2+^ may lead to different characteristics of SWS firing pattern. Alternatively, these hyperpolarization pathways may serve redundantly for the induction of the down state, leading to the robustness of the production of SWS firing pattern. In this study we investigated the possible role of Na^+^ dependent hyperpolarization pathway in the induction of SWS firing pattern. We modified the previous AN and SAN models to incorporate the changes in the kinetics of the voltage-gated Na^+^ channel and focused on the quantitative difference between Na^+^-dependent hyperpolarization pathway and Ca^2+^-dependent hyperpolarization pathway. Our results demonstrate that Na^+^ dependent hyperpolarization pathway is one of the candidate mechanisms that induce the down state of SWS firing pattern.

## RESULTS

### Construction of the NAN model by incorporating UNaV channel

Tatsuki *et al*., 2016 predicted that Ca^2+^-dependent hyperpolarization pathway contributes to the generation of SWS firing pattern with AN model^7^. In this model, intracellular Ca^2+^ gradually accumulates during the up state, and this accumulation activates KCa channels to induce the transition from the up state to the down state. During the down state, intracellular Ca^2+^ gradually diminishes over time because of Ca^2+^ pumps/exchangers action. However, the AN model as well as subsequent simplified SAN model^9^ did not consider the role of Na^+^ dynamics that may be capable of inducing transition from the up state to the down state. A possible mechanism for the Na^+^-dependent down state induction could be: activation of KNa currents during the up state to induce hyperpolarization, and subsequent decrease of intracellular Na^+^ by Na^+^ pumps/exchangers. Inward Na^+^ currents would be mainly mediated by voltage-gated Na^+^ channels and leak Na^+^ channels.

To test whether such Na^+^-dependent mechanism can be the underlying mechanism for SWS firing pattern, we constructed an alternative SAN model considering the role of the Na^+^ dynamics, called Na^+^-centered averaged-neuron model (NAN model) (**Figure 1A**). This model leverages mean-field approximations for a population of neurons to extract characteristic dynamics independently of specific neural circuits. Na^+^ dynamics was implemented by voltage-gated Na^+^ channels, leak Na^+^ channels and Na^+^ pumps/exchangers. We also considered the role of the KNa channels (**Figure 1A**).

**Figure 1.**
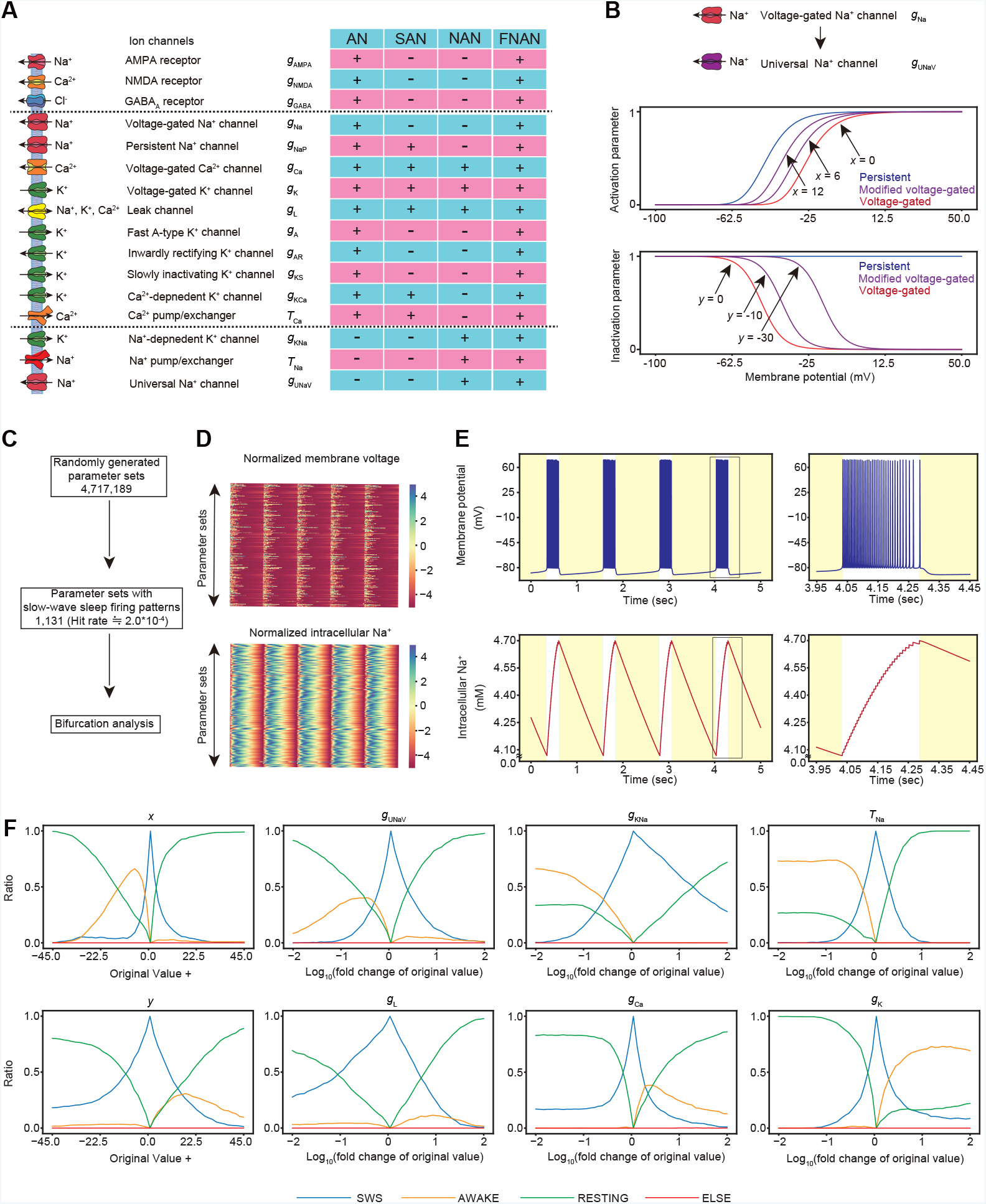
Construction of the NAN model by incorporating UNaV channel. **(A)** Schematic diagram of the AN, SAN, NAN and FNAN models. Some of the channels and pumps included in the AN model^7^ are included in the other models. In the NAN and FNAN models, KNa channels, UNaV channels, and Na^+^ pumps/exchangers are newly incorporated. **(B)** Changes in the activation/inactivation curve of UNaV channels alter the persistency of UNaV currents. (Left panel) Blue curve denotes the activation curve of persistent Na^+^ channels, while purple curves denote the activation curve of modified voltage-gated Na^+^ channels (Universal Na^+^ channels). Red curve in the left figure denotes the activation curve of voltage-gated Na^+^ channels with typical voltage sensitivity. When the value of parameter *x* is increased, purple curve shifts leftward (i.e., increased persistency). (Right panel) Blue, purple, and red curves denote the inactivation curve of corresponding channels as shown in left panel. When the value of parameter *y* is decreased, purple curve shifts rightward (i.e., increased persistency). Mathematical formulas for these channels are taken from Tatsuki et al., 2016^7^, which is based on Compte et al., 2003^10^. **(C)** Workflow of random parameter search. 1,131 parameter sets of the NAN model with SWS firing patterns were obtained with random parameter search. The SWS parameter sets were subsequently used in the bifurcation analysis. **(D)** The normalized membrane potential and intracellular Na^+^ are shown for all the 1,131 SWS parameter sets. **(E)** Representative SWS firing patterns and intracellular Na^+^ concentration of the NAN model. Right panels are magnified images. **(F)** The results of the bifurcation analysis with all parameters in the NAN model. Each panel explains the result of the bifurcation analysis for indicated parameter. Each channel conductance or time constant was gradually changed within the range from 10^−3^ to 10 times its original value. The *x* and *y* parameters were changed within the range from minus 45.0 to plus 45.0 from the original value. Vertical axis shows the ratio of parameter sets showing the indicated firing pattern (SWS, AWAKE, RESTING or other pattern). Criteria for the classification are based on the previous study^7^ (see **METHOD DETAILS**).

Persistent Na^+^ currents are incorporated in both AN and SAN models^7,9^ and the inactivation and activation curves of the persistent Na^+^ currents are illustrated in **Figure 1B**. As for the Na^+^ current, here we consider voltage-gated Na^+^ channels, which can be described as:

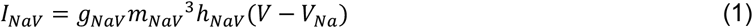

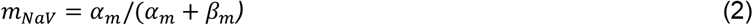

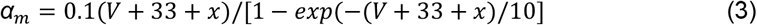

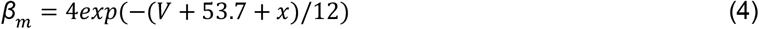

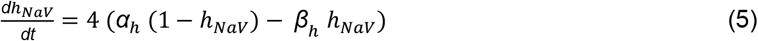

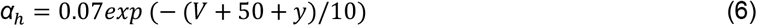

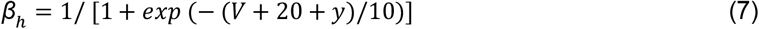

where *g*_NaV_ is the conductance of the voltage-gated Na^+^ channels, m_NaV_ is the activation parameter of the voltage-gated Na^+^ channels, h_NaV_ is the inactivation parameter of the voltage-gated Na^+^ channels, V_Na_ is the reversal potential for of voltage-gated Na^+^ channels, V is the membrane voltage, and t is time.

In these equations, the activation parameter m_NaV_ denotes the voltage sensitivity for the channels’ activation. The inactivation parameter h_NaV_ denotes the voltage sensitivity for the channels’ inactivation. It has been shown that these voltage sensitivities (i.e., m_NaV_ and/or h_NaV_) can be dynamically modulated by post-translational modifications. Thompson et al., 2017 reported that phosphorylation of SCN2a1 by CaMK2 resulted in the right shift of the inactivation curve^17^ and Daskal and Lotan reported that phosphorylation of the rat brain Na^+^ channels by PKC resulted in the right shift of the activation curve^18^. Failure of inactivation can lead to Na^+^ currents that have some degree of persistency against voltage dynamics (here we call this current as “persistent-like” Na^+^ currents). In our model scheme, this “persistent-like” Na^+^ currents can be described either by increasing the parameter *x* or decreasing the parameter *y* in Equation (3-7). An increase in the *x* results in the upregulation of currents mediated by the channels because *x* represent the shift of the activation curve and the leftward shift allows the channels to be activated at a lower membrane voltage, while a decrease in *y* results in the upregulation of currents mediated by the channels (**Figure 1B**) because *y* represent the shift of the inactivation curve and the rightward shift allows the channels to remain active at a lower membrane voltage. Thus, either of increasing *x* or decreasing *y* lead to the channels’ activity similar to that of persistent Na^+^ channels (**Figure 1B**). Note that if *x* = *y* = 0, the equations represent voltage-gated Na^+^ channels without such voltage-sensitivity modifications and are used in the AN and other models^7,9^. To distinguish these “normal” setting for the voltage-gated Na^+^ channels, we will call the voltage-gated Na^+^ channels with parameter *x* and *y* (i.e., *x* ≠ 0 and *y* ≠ 0) as universal voltage-gated Na^+^ (UNaV) channels.

The dependence of activation parameter of KNa channels on intracellular Na+ concentration was mimicked by fitting the data reported previously^12^. On the other hand, KCa channels and Ca^2+^ pumps/exchangers involved in the SAN model were not considered in the NAN model assuming that the roles of Ca^2+^-dependent K^+^ currents can be replaced by Na+-dependent K^+^ currents. The other receptors and channels found in the SAN model were incorporated in the NAN model because they are necessary to induce a bursting pattern and transition between AWAKE and SWS pattern^9^ (**Figure 1A**). Na^+^ pumps/exchangers were introduced as a linearized pump, where the speed of Na^+^ uptake follows sodium current and intracellular Na^+^ concentration. In short, the differences between the SAN and NAN model are the presence of UNaV channels, KNa channels and Na^+^ pumps/exchangers, and the absence of KCa channels and persistent Na^+^ channels.

To find parameter sets that produce SWS firing pattern by the NAN model, we comprehensively searched for parameters describing the conductance of ion channels, time constant for Na^+^ efflux, the parameter *x* and *y* for UNaV channels (**Figure 1C**). Among the 4,000,000 randomly generated parameter sets, 3.0 × 10^−4^ % (∼1,000) of them showed SWS firing pattern, suggesting that the essential roles of KCa currents in generating the SWS firing pattern can be replaced by KNa currents. SWS firing pattern exhibited intracellular Na^+^ waves that were coupled with the membrane potential (**Figure 1D, 1E**).

To explore the potential roles of each parameter in the generation of SWS firing pattern, we performed bifurcation analysis, where each channel conductance or time constant was gradually changed within the range from 10^−3^ to 10 times its value. The *x* and *y* parameters were changed within the range from minus 45.0 to plus 45.0 from the original value (**Figure 1F**).

**Figure 1F and Figure S1A** show bifurcation diagrams for each parameter in the NAN model. Increase of *x* resulted in the transition from the AWAKE firing pattern to the SWS firing pattern (see yellow and cyan curve). This is consistent with the anticipated role of Na^+^ dependent hyperpolarization pathway in the induction of the down state. Increase of *x* leads to effective Na^+^ accumulation at lower membrane voltage, which can activate KNa channels to induce the hyperpolarization for the down state. Similarly, increased Na^+^ accumulation by lower *y* or higher conductance of UNaV channels (*g*_UNaV_) also leads to the transition from AWAKE firing pattern to the SWS firing pattern, further support the role of Na^+^ dependent hyperpolarization pathway in the induction of the down state. Interestingly, parameter *x* may have a strict constraint for the induction of SWS firing pattern. This is evident from the narrow range of SWS pattern (cyan curve) observed in the *x* bifurcation diagram compared to those in *y* and *g*_UNaV_ (**Figure 1F**). The strict constraint for the parameter *x* is also evident from the distribution of parameters producing SWS firing pattern (**Figure S1B**). The distribution also indicate that the value of *x* should strictly be a positive value, meaning that lower voltage threshold for the activation curve of UNaV channels is necessary for the induction of SWS firing pattern in the NAN model. On the other hand, distribution of parameter *y* also suggests the mild constrain of this parameter for producing SWS firing pattern (**Figure S1B**).

Importance of intracellular Na^+^ level for the induction of down state in the NAN model is also supported by the observation that increase of the conductance of the KNa channels (*g*_KNa_) and the time constant of Na^+^ pumps/exchangers (*τ*_Na_) resulted in the transition from AWAKE firing pattern to SWS firing pattern (see yellow and cyan curve). This is consistent with the anticipated role of Na^+^ dependent hyperpolarization pathway driven by *g*_KNa_. Increase of *τ*_Na_ should lead to effective accumulation of intracellular Na^+^. Interestingly, *g*_KNa_ may have a loose constraint for the induction of SWS firing pattern.

On the other hand, increase of the conductance of the voltage-gated Ca^2+^ channels (*g*_Ca_) or the conductance of the voltage-gated K^+^ channels (*g*_K_) resulted in the transition from SWS firing pattern to AWAKE firing pattern. The NAN model does not involve the KCa channels and thus the main role of the voltage-gated Ca^2+^ channels in the NAN model may be to upregulate the excitability of neurons. The roles of voltage-gated K^+^ channels for the induction of AWAKE firing pattern can be estimated by a detailed investigation of intracellular Na^+^ level: under the higher *g*_K_, intracellular Na^+^ level is not accumulated even in the up state, indicating that Na^+^ influx is attenuated (**Figure S1C**).

We also note that changes in the conductance of the leak channels (*g*_L_) had less effect in the SWS/AWAKE transition, indicating that leak channels are not important in the generation of the SWS firing pattern in the NAN model.

The NAN model does not include ion channels depending on the intracellular Ca^2+^ (e.g., KCa channels). However, we still keep the voltage-gated Ca^2+^ channels in the NAN model because of the following reasons. We also tested different version of the NAN model without voltage-gated Ca^2+^ channels (NAN model with *g*_Ca_ = 0, **Figure S2A**). This model indeed has a capability to produce the SWS firing pattern. The distribution also indicate that the value of *x* should strictly be a positive value, meaning that lower voltage threshold for the activation curve of UNaV channels is necessary for the induction of SWS firing pattern in this model (**Figure S2B**). However, the bifurcation diagram of the conductance of the UNaV channels (*g*_UNaV_) indicate that either increase and decrease of the conductance lead to the transition from SWS to AWAKE firing pattern (compare *g*_UNaV_ panel in **Figure S2C and Figure 1F**). In short, the role of UNaV channels for SWS firing pattern is not clear. This is probably because Na^+^ currents inherently has two roles: facilitator of depolarizing and also facilitator of hyperpolarizing induced by Na^+^-dependent K^+^ currents. On the other hand, voltage-gated Ca^2+^ channels in the NAN model have only a role for the induction of depolarization. Thus, the presence of voltage-gated Ca^2+^ channels (i.e., original NAN model in **Figure 1A**) may support the role of UNaV channels as the facilitator of hyperpolarization by activating the Na^+^-dependent K^+^ currents, and the clear distinction of SWS/AWAKE transition in the *g*_UNaV_ bifurcation is appeared in the original NAN model.

### Accumulation of Na^+^ drives the transition from the up state to the down state

We investigated the mechanism of SWS firing pattern by analyzing the ODEs trajectory around the transition from the up state to the down state (**Figure 2A-B**). **Figure 2C-D** show that action potential during the up state is initiated by UNaV channels and continued by voltage-gated Ca^2+^ channels. The action potential is ceased by the activation of voltage-gated K^+^ channels and KNa channels. During the down state, most of the hyperpolarization currents are mediated by KNa current. This is in contrast to what the AN and SAN models observed showing the importance of KCa currents for the hyperpolarization during the down state. In the NAN model, leak K currents also constitute the hyperpolarization current, consistent with the SAN model. **Figure 2E-F** show that intracellular Na^+^ rises cumulatively during depolarization phase (i.e., the up state) and then continuously decreases after the onset of the down phase. The increase and decrease of intracellular Na^+^ concentration can be explained by the altered ratio of overall activity between UNaV/Leak Na^+^ channels(IN) and Na^+^ pump (OUT) (**Figure 2F**).

**Figure 2.**
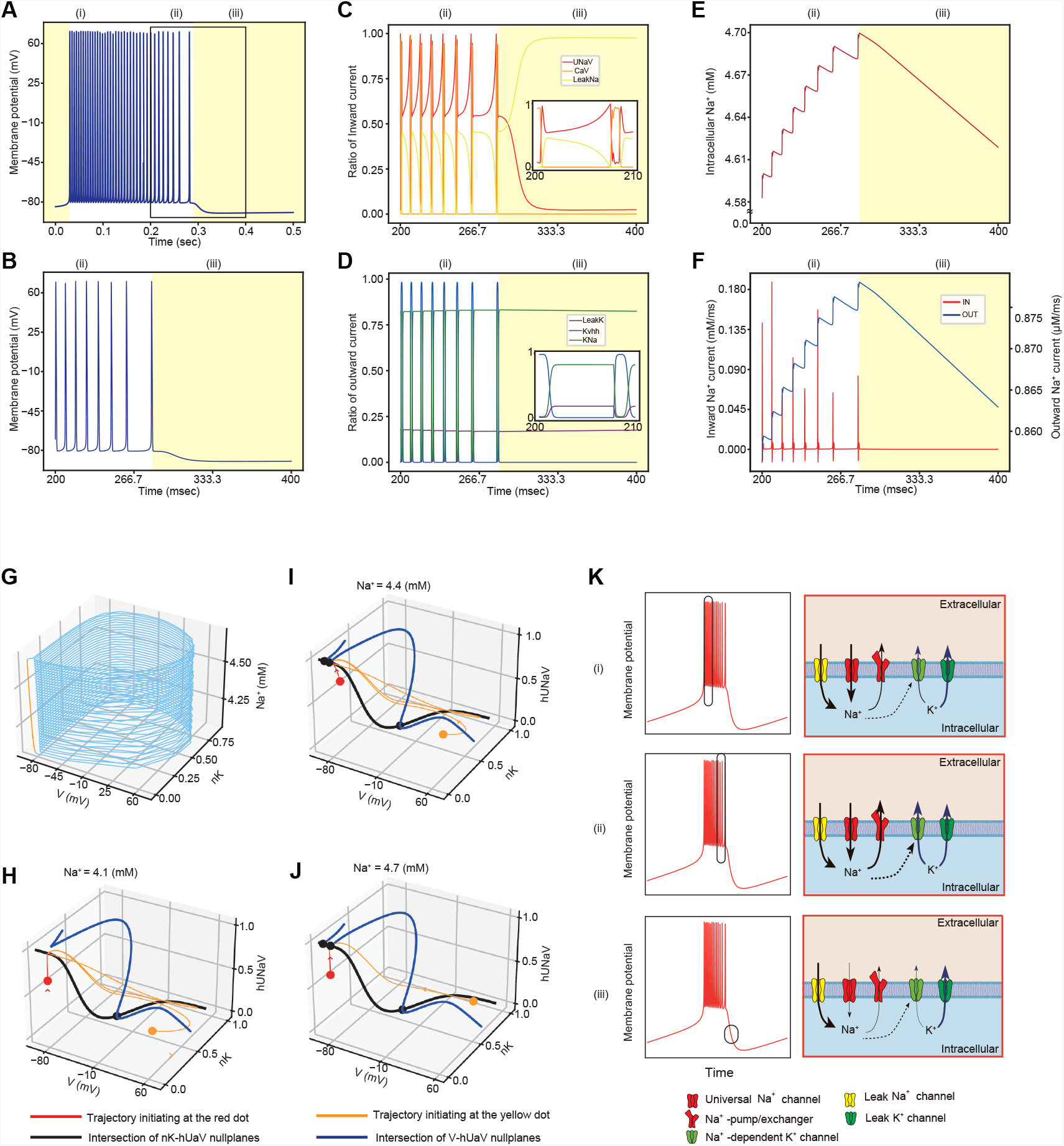
Accumulation of Na^+^ drives the transition from the up state to the down state. (A-B) Representative time course of membrane voltage in the NAN model. The parameter set for this graph is the same parameter set for **Figure 1E**. Yellow areas denote the down state whereas white areas denote the up state. The area surrounded by the black square indicates the place where it is magnified in panels. (B)is the Magnified image of (A). (C-F) The time in **Figure 2C-F** is the same as the time in **Figure 2B**. (C) Time course of inward currents. “UNaV” denotes the currents with UNaV channels, “CaV” denotes the currents with voltage-gated Ca^2+^ channels and “LeakNa” denotes currents with leak Na^+^ channels. Bottom-right figure is the magnification of the time courses of these currents. (D) Time course of outward currents. “LeakK” denotes the currents with leak K^+^ channels, “Kvhh” denotes the currents with voltage-gated K^+^ channels, “KNa” denotes the currents with KNa channels. Bottom-right figure is the further magnification of the time courses of these currents. (E) Time course of intracellular Na^+^. (F) Time course of the inward/outward Na^+^ currents. (G) Trajectory of the NAN model plotted on 3-variables space: membrane voltage (V), activation parameter of the voltage-gated K^+^ channel (nK), and intracellular Na^+^. The blue line denotes the trajectory corresponding to the up state, whereas the orange line denotes the trajectory corresponding to the down state. (H-J) Intersection of V, inactivation parameter of the universal Na^+^ channel (hUNaV) nullplanes is represented by blue curve, whereas Intersection of nK, hUNaV nullplanes is represented by black curve. Red or yellow dot denotes the initial condition of the integration with ODEs. Black dots denote the fixed point(s) of the system. To reduce the number of variables from four to three, Na^+^ is fixed at indicated values (4.1 mM (H), 4.4 mM (I), or 4.7 mM (J)). (K) Schematic representation of the mechanisms generating SWS firing pattern with Na^+^-dependent hyperpolarization pathway. (i) During the up state, there is transient Na^+^ inward currents, and Na^+^ is pumped out by Na^+^ pumps. (ii) Gradually intracellular Na^+^ accumulates and up states is terminated because KNa currents are dominant. (iii) During the down state, intracellular K^+^ exits the cell mainly through KNa channels. Note that (i)-(iii) shown in **Figure 2A-F** correspond to the scheme (i)-(iii).

To analyze the mechanism behind the induction of the down state, we plotted the trajectory of SWS firing pattern in the phase space (**Figure 2G**). The NAN model has four variables in the ODEs (i.e., V, the membrane potential; nK, the dimensionless quantity associated with the activation of the voltage-gated K^+^ channels; hUNaV; the dimensionless quantity associated with the inactivation of UNaV channels, and Na^+^; the intracellular Na^+^ concentration). The coiled part (green trajectory) and the straight part (yellow trajectory) correspond to the up state and the down state, respectively. **Figure 2H-J** plotted the intersection of nullplanes at fixed Na^+^ concentrations. As the nullplane is a set of the points in which the first derivative of a certain direction is zero, a system on the intersection of two nullplanes only evolves to the remaining variable: for example, on the intersection of V-nK nullplanes, only the hUNaV variable shall evolve. Therefore, the nullplane analysis enables us to understand the approximate dynamics of the ODEs.

At the low Na^+^ concentration (4.1 mM), the trajectory converged to the stable limit cycle (**Figure 2H**), which corresponded to the up state of the SWS firing pattern. At the middle Na^+^ concentration (4.4 mM), there were at least two stable states: a stable point and a stable limit cycle, which correspond to the down state and the up state, respectively (**Figure 2I**). At the high Na^+^ concentration (4.7 mM), the stable limit cycle does not exist, and the trajectory converged to the stable point corresponding to the down state (**Figure 2J**). In summary, the phase planes of the NAN model indicate that the alteration between the up and the down state is associated with a Na^+^ concentration-dependent transition of stable states (**Figure 2K)**.

### Parameter *x* is important for controlling several features of wave pattern in the NAN model

Next we investigated how several features of wave patterns produced in the NAN model are regulated by each parameter. Each SWS parameter found in random parameter search (**Figure 1C**) was slightly decreased or increased, and the effect of slight change on SWS firing pattern was quantified. To characterize the SWS firing pattern, we focused on the duration of the up state and the down state, oscillation period (defined as the duration of the up state + the down state), amplitude of intracellular Na^+^ oscillation, inter-spike interval (ISI) during the up state. **Figure 3A-G** showed that parameter *x* elongated the period of the down state, oscillation period, Na^+^ oscillation amplitude, and shortened ISI. This suggests that *x* plays a major role in the determination of SWS oscillation and Na^+^ dynamics in this model.

**Figure 3.**
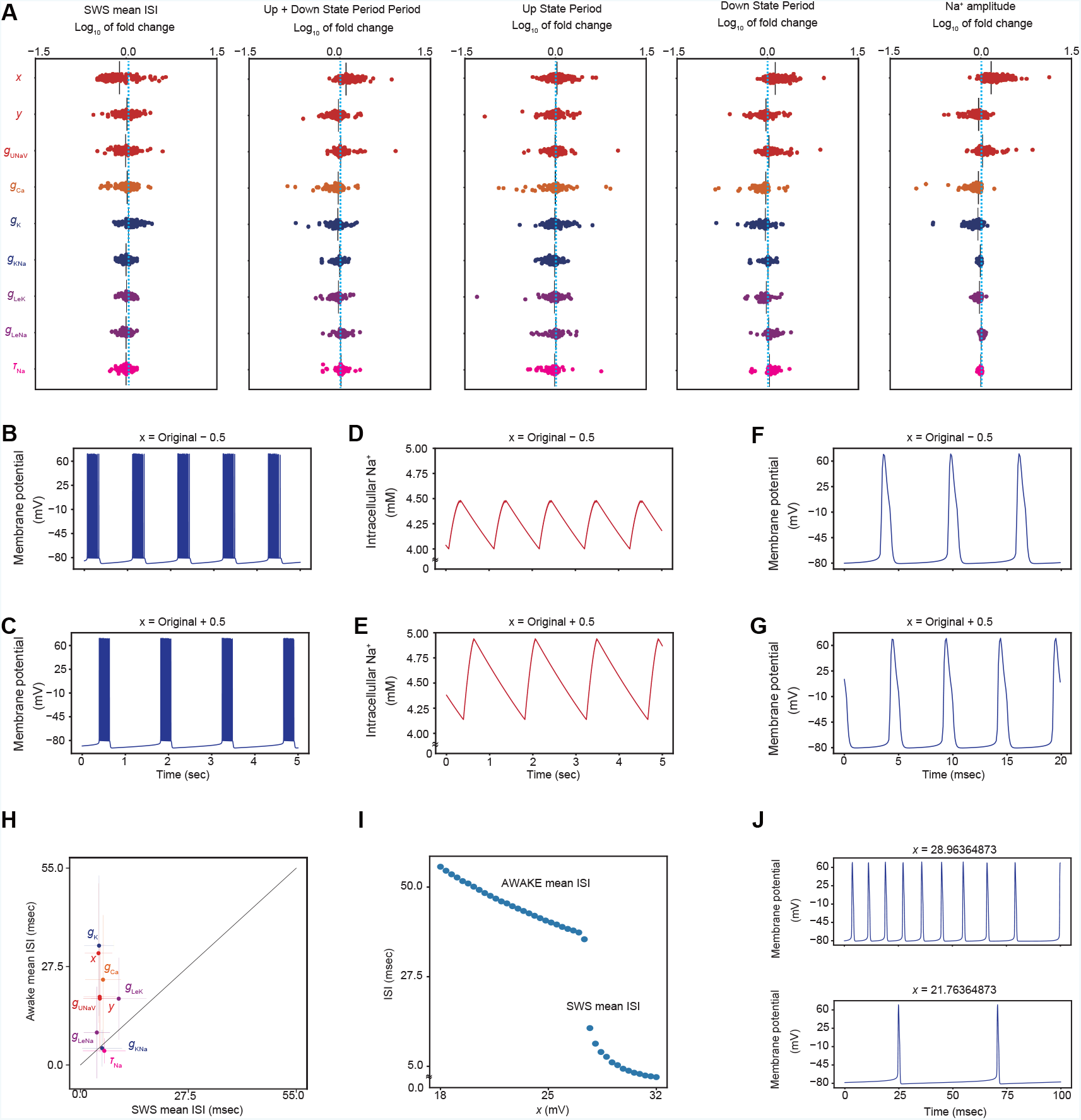
Activation parameter of UNaV is important for controlling several features of wave pattern in the NAN model. (A) The effect of slight upregulation of the value of each parameter alter the characteristics of SWS firing pattern. All parameter sets yielding SWS firing pattern are used in the analysis. For parameter *x* and *y*, characteristics of SWS firing pattern is calculated when the value of *x, y* is (i): 0.5 lower than the original value (ii): 0.5 higher than the original value. For the other parameters, characteristics of SWS firing pattern is calculated when the value is (i): 0.975 times of the original value (ii): 1.025 times of the original value. The value calculated in (ii) is divided by the value calculated in (i). Blue dotted line shows Log_10_ of fold change = 0 (i.e. perturbation does not alter each characteristic of SWS firing pattern). (B, C) Representative trace of the firing pattern when the value of *x* is 0.5 lower than the original value (B) and 0.5 higher than the original value (C). (D, E) Representative trace of the intracellular Na^+^ oscillation when the value of x is 0.5 lower than the original value (D) and 0.5 higher than the original value (E). (F, G) Representative trace showing the ISI when the value of x is 0.5 lower than the original value (F) and 0.5 higher than the original value (G). (H) The trends of ISI in SWS and AWAKE firing patterns. To induce AWAKE firing pattern for the majority of parameter sets with SWS firing pattern, the amount of perturbation was selected based on the bifurcation analysis in **Figure 1F** so that the induction of AWAKE firing pattern is the most effective (*x*; -7.2, *y*; +16.2, *g*_K_; ×10^1.52^, *g*_Ca_; ×10^0.4^, *g*_LeK_; ×10^−2.0^, *g*_LeNa_; ×10^0.88^, *g*_UNaV_; ×10^−0.48^, *g*_KNa_; ×10^−2.0^, *τ*_Na_; ×10^−0.96^). Parameter sets which shows AWAKE firing pattern when each parameter is changes as above are selected for each parameter’s analysis. Mean ISI of the firing pattern is calculated for each parameter set, and dots with color denotes the average of mean ISI among parameter sets. (I) Representative change of mean ISI when the value of parameter *x* is gradually changed in the representative parameter set. (J, K) Representative trace showing ISI when the firing pattern is SWS firing pattern (J) and AWAKE firing pattern (K). The transition from SWS to AWAKE is induced by lowering the value of parameter *x*.

It has been shown that ISI during the up state of SWS firing pattern tend to be smaller compared to the ISI during the wake time^19,20^. Similar trend can be observed when the bifurcation from the SWS firing pattern to AWAKE was induced by changing the parameters *x, y, g*_K_, *g*_Ca_, *g*_UNaV_. On the other hand, such reduced ISI in the up state was hardly observed when the bifurcation was induced by changing *g*_KNa_, *τ*_Na_ (**Figure 3H-J**). Parameters *x, y, g*_K_, *g*_Ca_ and *g*_UNaV_, but not *g*_KNa_ and *τ*_Na_, are all related to the ion channels whose activation is directly regulated by membrane potential. This result suggests that bifurcation of SWS to AWAKE firing pattern *in vivo* neuron is related to changes in voltage-related channel regulation.

### Na^+^-dependent hyperpolarization pathway can also be implemented by Na^+^/K^+^ ATPase

In the NAN model, Na^+^ pumps/exchangers are described by Na^+^ pumps with a linear kinetics. In the actual neurons, however, majority of Na^+^ efflux is mediated by Na^+^/K^+^ ATPases that exchange 2 K^+^ and 3 Na^+^ ions coupled with ATP hydrolysis^21^. To investigate the possible contribution of Na^+^/K^+^ ATPase as a molecular nature for the export of Na^+^ from the neurons, we created an alternative model (**Figure 4A**). We speculated that hyperpolarization driven by Na^+^/K^+^ ATPases could be the main source of hyperpolarization currents inducing the down state. To focus on such a role for the Na^+^/K^+^ ATPases, the revised model with Na^+^/K^+^ ATPases does not include KNa channels, which support the generation of the the down state in the NAN model as described above. Among the 50,000,000 randomly generated parameter sets, 2.0 × 10^−5^ % (∼1,000) of them showed SWS firing pattern in the revised NAN model, suggesting that the essential roles of KNa currents in generating the SWS firing pattern can be replaced by Na^+^/K^+^ ATPases currents. Intracellular Na^+^ concentration oscillates with the membrane potential (**Figure 4B, 4C**). **Figure 4D and Figure S4A** show bifurcation diagram in the revised NAN model. Decrease of the conductance of the Na^+^/K^+^ ATPases (*g*_NaK_) resulted in the transition from the SWS firing pattern to the AWAKE firing pattern, supporting the importance of Na^+^/K^+^ ATPases mediated hyperpolarization currents for the induction of the down state. **Figure 4E** and **Figure S4B** show the distribution of each parameter. The distribution of *g*_NaK_ found in the SWS parameter sets is restricted in the larger *g*_NaK_ (**Figure 4E**), suggesting that sufficiently large *g*_NaK_ is required to generate SWS firing pattern.

**Figure 4.**
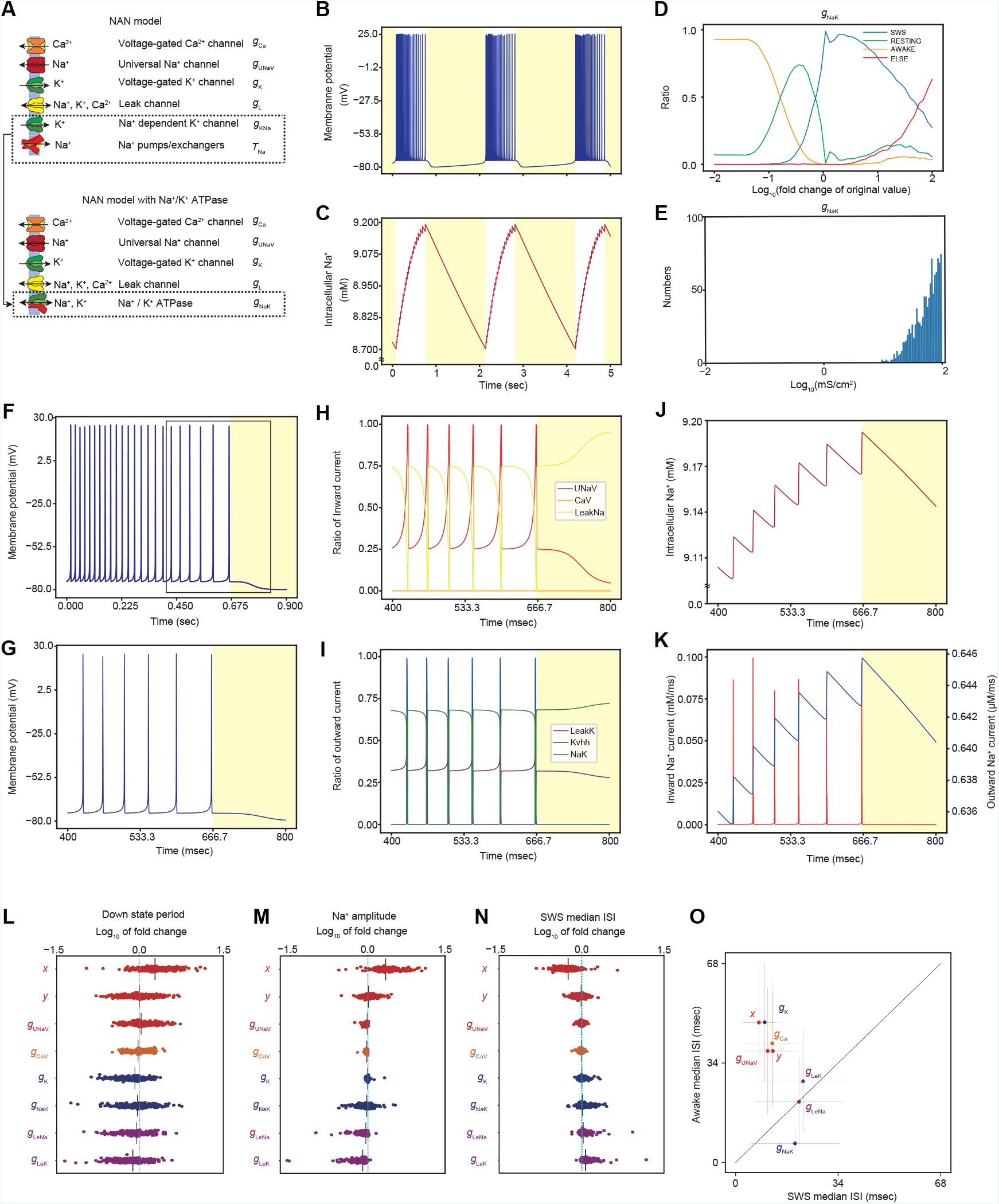
Na^+^-dependent hyperpolarization pathway can also be implemented by Na^+^/K^+^ ATPase. (A) Schematic diagram of the NAN model with Na^+^/K^+^ ATPases. (B, C) Representative SWS firing pattern and intracellular Na^+^ concentration of the NAN model with Na^+^/K^+^ ATPases. (D) Results of the bifurcation analysis with all parameter sets by changing the value of parameter *g*_NaK_. Classification criteria of SWS, AWAKE, RESTING, ELSE follows that of Tatsuki et al., 2016^7^. (E) The distributions of the value of the conductance of Na^+^/K^+^ ATPases in all parameter sets yielding SWS firing pattern in the NAN model with Na^+^/K^+^ ATPases. (F-G) Representative time course of membrane voltage in the NAN model with Na^+^/K^+^ ATPases. The area surrounded by the black square indicates the place where it is magnified in the subsequent imaging. (G) is the the magnified image of (F) in the area surrounded by the black square. (H-K) The time in **Figure 4H-K** is the same as the time in **Figure 4G.** (H) Time course of inward currents. “UNaV” denotes the currents with UNaV channels, “CaV” denotes the currents with voltage-gated Ca^2+^ channels and “LeakNa” denotes currents with leak Na^+^ channels. (I) Time course of outward currents. “LeakK” denotes the currents with leak K^+^ channels, “Kvhh” denotes the currents with voltage-gated K^+^ channels, “NaK” denotes the currents with Na^+^/K^+^ ATPases. (J) Time course of intracellular Na^+^. (K) Time course of the inward/outward Na^+^ currents. (L, M, N) The effect of slight upregulation of the value of each parameter alter the characteristics of SWS firing pattern. All parameter sets yielding SWS firing pattern are used in the analysis. For parameter *x* and *y*, characteristics of SWS firing pattern is calculated when the value of *x, y* is (i): 0.5 lower than the original value (ii): 0.5 higher than the original value. For the other parameters, characteristics of SWS firing pattern is calculated when the value is (i): 0.975 times of the original value (ii): 1.025 times of the original value. The value calculated in (ii) is divided by the value calculated in (i). Blue dotted line shows Log_10_ of fold change = 0 (i.e. perturbation does not alter each characteristic of SWS firing pattern). (O) The trends of ISI in SWS and AWAKE firing pattern. To induce AWAKE firing pattern for the majority of parameter sets with SWS firing pattern, the amount of perturbation was selected based on the bifurcation analysis in **Figure 4D, Figure S4D** so that the induction of AWAKE firing pattern is the most effective (*x*; -3.6, *y*; +7.0, *g*_K_; ×10^0.48^, *g*_Ca_; ×10^0.32^, *g*_LeK_; ×10^−1.84^, *g*_LeNa_; ×10^0.48^, *g*_UNaV_; ×10^−0.4^, *g*_NaK_; × 10^−1.44^). Parameter sets which shows AWAKE firing pattern when each parameter is changes as above are selected for each parameter’s analysis. Mean ISI of the firing pattern is calculated for each parameter set, and dots with color denotes the average of mean ISI among parameter sets.

**Figure 4F-I** show that action potential during the up state is initiated by UNaV channels. The action potential is ceased by the activation of voltage-gated K^+^ channels and Na^+^/K^+^ ATPases. During the down state, ∼75 % of the hyperpolarization current is mediated by Na^+^/K^+^ ATPases currents. The rest of ∼25 % hyperpolarization current is mediated by leak K^+^ currents, consistent with the SAN model^9^. **Figure 4J-K** show that intracellular Na^+^ rises cumulatively during depolarization phase (i.e., the up state) and then continuously decreases after the onset of the down phase. The increase and decrease of intracellular Na^+^ concentration can be explained by the altered ratio of overall activity between UNaV/Leak Na (IN) and Na^+^/K^+^ ATPase (OUT) (**Figure 4K**).

The role of parameter *x* for shaping the SWS-firing pattern is consistent between the NAN model and the revised NAN model: increase in parameter *x* elongated down state duration, Na^+^ oscillation amplitude, and decreased ISI in the up state during SWS firing pattern (**Figure 4L-O and Figure S4C**). Similar to the case of *g*_NaK_, perturbation on *g*_NaK_ also resulted in the decrease in the ISI across the transition from SWS to AWAKE state, which is opposite direction observed in the tendency of ISI transition in vivo, suggesting that other factors such as *x* might be responsible for the transition from SWS to AWAKE in vivo.

### The Full-NAN model reveals the importance of Na^+^-dependent hyperpolarization pathway

To evaluate the role of Na^+^-dependent hyperpolarization pathway in the presence of multiple pathways, some of which are not explicitly described in NAN (and SAN) model, we created a Full-NAN (FNAN) model. The FNAN model includes all channels/pumps in the original AN model^7^ as well as the NAN model. This model consists of three neurotransmitter receptors and 13 ion channels and pumps (**see Figure 1A**). Among the ∼24,000,000 randomly generated parameter sets, 8.9 × 10^−5^ % (∼2,000) of them showed SWS firing pattern. To ask which ion channels/pumps play an essential role for the generation of SWS firing pattern, we next performed “knockout experiments”, where we removed the contribution of each ion channels/pumps. Removal of GABAR currents did not disrupt SWS firing pattern in a majority of parameter sets (**Figure S5A**). We thus decided to exclude GABAR in the subsequent analysis. Knockout of KNa channels led to the transition from the SWS firing pattern to the AWAKE firing pattern in a majority of parameter sets, which otherwise produce SWS firing patterns. On the other hand, knockout of KCa channels did not alter the firing pattern in most cases (**Figure 5A**), indicating that in a majority of parameter sets in the FNAN model, Na^+^-dependent hyperpolarization (driven by KNa channels) is more dominant than Ca^2+^-dependent hyperpolarization (driven by KCa channels) in inducing the down state. Indeed, only a few parameter sets showed AWAKE firing pattern when the conductance of KCa channels is set to zero, and among these 40 parameter sets, 31 parameter sets also showed AWAKE firing pattern by the knockout of KNa channels (**Figure S5B**). In other words, KCa channels are only fully responsible for the induction of SWS firing pattern for only the remaining 9 out of 1,444 parameter sets. The prominent role of KNa channels or Na^+^-dependent hyperpolarization pathway is further supported by the representative trace of the FNAN model. **Figure S5C** shows that peak and trough of Na^+^ dynamics are correlated to the beginning and ending of the down state, whereas those of Ca^2+^ dynamics do not show consistent increase or decrease during the up or down state, respectively.

**Figure 5.**
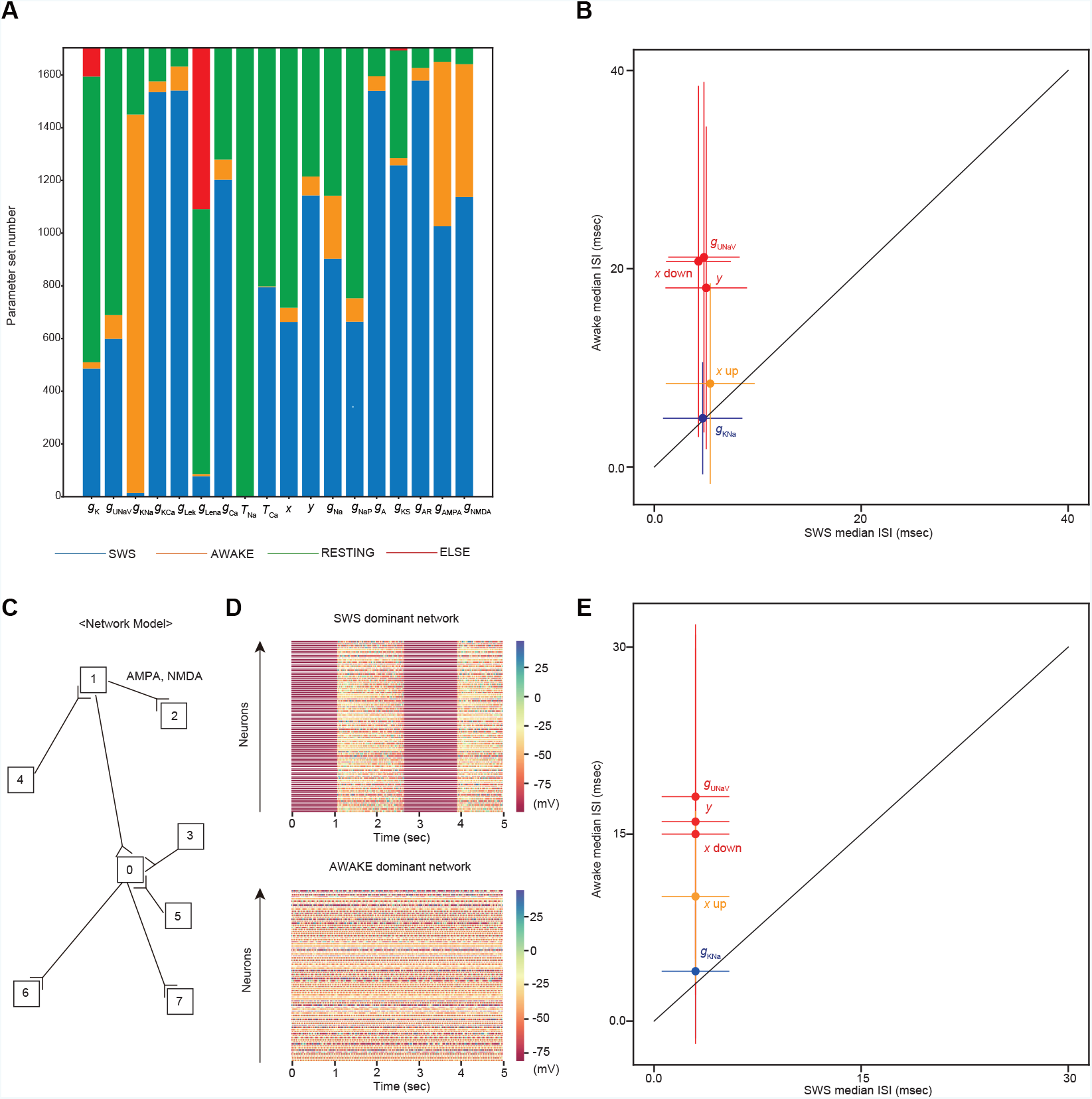
The Full-NAN model reveals the importance of Na^+^-dependent hyperpolarization pathway. (A) The result of knockout analysis with all parameter sets showing SWS firing pattern. When the contribution of a particular element is set to zero, the number of parameter sets showing AWAKE firing pattern is shown by orange, and the amount of parameter sets showing RESTING firing pattern is shown by green and the amount of parameter sets which are excluded from the analysis (ELSE) because of their aberrant firing pattern is shown by red, and the amount of parameter sets showing SWS firing pattern is shown by blue. Classification criteria of SWS, AWAKE, RESTING, ELSE follows that of Tatsuki et al., 2016^7^. (B) The trends of ISI in SWS and AWAKE firing pattern. To induce AWAKE firing pattern for the majority of parameter sets with SWS firing pattern, the amount of perturbation was selected based on the bifurcation analysis in **Figure S5A** so that the induction of AWAKE firing pattern is the most effective (*x*; +12.6, *y*; +14.4, *g*_UNaV_; × 10^−0.72^, *g*_KNa_; × 10^−2.0^). Mean ISI of the firing pattern is calculated for each parameter set, and dots with color denotes the average of mean ISI among parameter sets. (C) Construction of network model. The network is consisted of excitatory neurons. Hence, neurons are connected with AMPA or NMDA-mediated synapses. (D) Representative trace of all neurons in the network model. The upper figure shows a SWS-dominant network, whereas the lower figure shows an AWAKE-dominant network, where the conductance of KNa channels is set to be 0.01 times of what it is in the left figure. (E) The trends of ISI in SWS and AWAKE firing patterns. ISI in SWS firing patterns in Figure 5F “normal” is compared with AWAKE firing pattern in Figure 5F “*g*_KNa_ down”, “*g*_UNaV_ down”, “y up”, “*x* down”, “*x* up”. Among the all parameter sets showing SWS firing pattern, changes of the firing pattern from SWS firing pattern to AWAKE firing pattern is seen when the value of the parameter is modified in that the majority of the parameter sets show AWAKE firing pattern instead of SWS firing pattern. The exact amount of perturbation given to each parameter is determined by the results of the bifurcation analysis in **Figure S5A** (*x* down; -12.6, *x* up; +12.6, *y*; +14.4, *g*_UNaV_; ×10^−0.72^, *g*_KNa_; ×10^−2.0^). Parameter sets which shows AWAKE firing pattern when each parameter is changes as above are selected for each parameter’s analysis. Mean ISI of the firing pattern is calculated for each parameter set, and dots with color denotes the mean ISI of mean ISI in parameter sets.

The role of UNaV channels, particularly for the parameter *x* is mostly conserved in the FNAN and NAN models. Similar to the case of the NAN model, shortening of ISI in the up state of SWS firing patten was observed when AWAKE was induced by changing the parameters *x, y, g*_UNaV_ (**Figure 5B**). Such trend was hardly observed when the bifurcation was induced by changing *g*_KNa_, *τ*_Na_ in the FNAN model, also consistent with the case of the NAN model.

Furthermore, the FNAN model obscures the consistent change of wave features (i.e. Down state period, Na^+^ amplitude, and ISI in the up state) upon the increase/decrease of parameter *x*. (**Figure S5D**; see also **Figure 3A**). This is possibly because the FNAN model includes the redundant sources of inward Na^+^ currents such as voltage-gated Na^+^ channels and persistent Na^+^ channels.

To investigate whether the role of Na^+^-dependent hyperpolarization pathway revealed so far requires the assumption in mean-field approximation of the model, we created a neuronal network with the FNAN model, in which neurons are explicitly connected with NMDA and AMPA-mediated synaptic connections without the mean-field approximation (**Figure 5C**). Note that we set *g*_GABA_ to be 0 because SWS firing pattern persists without GABAergic currents^2^. We simulated the behavior of network FNAN model with 100 representative parameter sets. **Figure 5D** shows a representative behavior of the network model showing SWS firing pattern and AWAKE firing pattern. We subsequently investigated whether the tendency of ISI observed in the FNAN/NAN models are also preserved in this network model (**Figure S6A**). Perturbations to each parameter are given so that the wave pattern changes from SWS to AWAKE. The amount of perturbation is determined based on the bifurcation diagram of each parameter (**Figure S6B**). Similar to the case of single cell FNAN model (**Figure 5B**), shortening of ISI in the up state of SWS firing patten was observed when AWAKE was induced by changing the parameter *x, y, g*_UNaV_ (**Figure 5E**). Such trend was hardly observed when the bifurcation was induced by changing *g*_KNa_ in the network FNAN model, which is also consistent with the case of single cell FNAN model.

To summarize, at least part of the role of Na^+^-dependent hyperpolarization pathway in the NAN model is conserved in the FNAN model regardless of the assumption of mean-field approximation.

## DISCUSSION

### Putative mechanism for SWS firing pattern with the NAN model

In this study, we developed the NAN model, which induces the down state through the activation of K^+^ currents triggered by intracellular Na^+^. The NAN model revealed that KNa channels induce the down state. The SWS firing pattern is generated through a series of events summarized as: (i) during the up state, Na^+^ enters mainly through voltage-gated Na^+^ channels and leak Na^+^ channels to increase the intracellular Na^+^ concentration. (ii) The transition from the up to the down state occurs when the intracellular Na^+^ concentration reaches a certain threshold to activate Na^+^-dependent K^+^ channels. (iii) during the down state, Na^+^ exits through Na^+^-pump/exchangers to decrease the intracellular Na^+^ concentration. In summary, intracellular Na^+^-dependent hyperpolarizing current drives the transition from the up state to the down state. Compte et al., 2003 indicated the importance of KNa channels by leveraging complex excitatory-inhibitory neuronal networks^10^. Our study showed the significance of KNa channels independent of such intricate connections and without the contribution of inhibitory neurons. The Na^+^-dependent hyperpolarization pathway may be implemented by several molecules other than KNa channels. For example, we demonstrated that Na^+^/K^+^ ATPases create a hyperpolarizing current dependent on the accumulation of the intracellular Na^+^ (**Figure 4**).

In the Na^+^-dependent hyperpolarization pathway, the activation of hyperpolarizing current should be initiated by neuronal activity-dependent manner. Increase in parameter *x* of the voltage-gated Na^+^ channels, which denotes the negative shift of the activation curve, results in the increase of neuronal activity. Thus, increased *x* leads to SWS firing pattern (through stimulating the neuronal activity-dependent hyperpolarization) and decreased ISI during up state (through stimulating the neuronal activity). The activation parameter *x* is also important in shaping the SWS firing patterns by modulating the Na^+^ oscillation period as well as Na^+^ oscillation’s amplitude (**Figure 3A**). Having the simulation results recapitulating the tendency of ISI upon the SWS/AWAKE alteration in vivo (i.e., ISI during the up state in SWS firing pattern tends to be shorter than ISI in the AWAKE firing pattern), parameter *x* is a candidate point to induce the changes between SWS firing AWAKE firing *in vivo* (**Figure 3H, 3I, 3J**). Activation/inactivation kinetics of voltage-gated Na^+^ channel can be altered by post-translational modification along with the sleep-wake cycle, because several kinases were reported to alter the activation/inactivation kinetics of voltage-gated Na^+^ channels^17,18^. Bruning *et al*., 2019 reported that phosphorylation level of voltage-gated Na^+^ channels is increased under light phase where the mice should have higher sleep needs^22^.

### Comparison with other mathematical models recapitulating up-down oscillation

The essence of the NAN model’s scheme is that during the up state the amount of intracellular Na^+^ which can induce the down state with the activation of KNa channels gradually rises, and during the down state the amount of intracellular Na^+^ gradually diminishes. The role of intracellular Na^+^ in the NAN model for the induction of the down state can be achieved by other molecules. Previous studies suggested that intracellular Ca^27,9,23-25^ may be the inducer of the down state. During the up state Ca^2+^ enters through voltage-gated Ca^2+^ channels and when it is accumulated, KCa channels are activated, inducing the down state. During the down state plasma membrane Ca^2+^ ATPases (Ca^2+^ pumps/exchangers) pump out intracellular Ca^2+^, reinitiating the up state. Another motif is that voltage dependent slowly activating K^+^ channels^26^, which is gradually activated during the up state and induces down state, and since during the down state membrane potential is hyperpolarized, so the channel is gradually inactivated.

The induction of the down state can be achieved by activity-dependent inactivation of depolarizing current. Voltage-dependent Na^+^ channels^27-29^, and Ca^2+^-dependent inactivation of Ca^2+^ channel^30^ are reported to be important for the induction of the down state. It would be important to investigate the relative contribution of activity-dependent induction of hyperpolarization and activity-dependent inactivation of depolarization for the generation of down state.

### Relevance of Na^+^-dependent hyperpolarization pathway with previous studies

Several lines of evidence support the involvement of Na^+^-dependent hyperpolarization pathway in the regulation of sleep. Bifurcation diagram in **Figure 1F** suggests that downregulation of KNa currents altered the firing pattern from SWS to AWAKE. One way to achieve the downregulation of KNa currents is to decrease intracellular K^+^ concentration compared to the extracellular K^+^ concentration. In other words, lower extracellular K^+^ concentration may lead to sleep. This is consistent with the previous study where Ding et al., 2016 showed that during sleep/anesthesia, extracellular K^+^ concentration falls^31^. Application of artificial cerebro-spinal fluid (ACSF) mimicking the cerebro-spinal fluid of the sleep state induced sleep in vivo, showing the causal effect of extracellular ion condition on sleep.

Figure 3. showed that the alteration of the activation parameter of voltage-gated Na^+^ channels is important for generating SWS firing pattern. Activation kinetics of voltage-gated Na^+^ channels may be dynamically regulated in sleep-wake cycle, and phosphorylation of voltage-gated Na^+^ channels is one candidate for the regulation. It is observed that phosphorylation level of one type of voltage-gated Na^+^ channels SCN2a1 is increased when sleep need is accumulated by the administration of MK-801 (NMDA receptor antagonist)^32^ or during light phase where mice typically undergo sleep^22^. Involvement of SCN2a in the sleep control is demonstrated by showing that Deficiency of Scn2a led to increased wakefulness and decreased NREM sleep duration^33^. Because the regulation of activation/inactivation kinetics of voltage-gated Na^+^ channel often include the phosphorylation of the channel^17,18^, it is possible that phosphorylation of SCN2a may lead to the left-shift of the activation parameter, leading to the induction of sleep as suggested by our model.

Bifurcation diagram in **Figure 4D** suggests that downregulation of Na^+^/K^+^ ATPases currents altered the firing pattern from SWS to AWAKE. This is supported by animal experiments: mice harboring an inactivating mutation in the neuron-specific Na^+^/K^+^ ATPase α3 subunit exhibit decreased sleep duration^34^. Also, injection of ouabain, an inhibitor of Na^+^/K^+^ ATPases, enhanced wakefulness in mice^35^.

### Limitations of the study

The importance of Na^+^-dependent hyperpolarization pathway for sleep need experimental validation. The NAN-model’s prediction about the changes in neuronal firing pattern upon the perturbation towards Na^+^-dependent hyperpolarization pathway (i.e., trend of bifurcation seen in **Figure 1F**, trend of up state/down state duration, ISI in **Figure 3** and **Figure 4**) would be of particular interest for the assessment.

The NAN model is based on Hodgkin-Huxley model, which has a limitation in modeling the dynamics of intracellular Na^+^/Ca^2+^ concentration because because the reversal potential is a fixed value. A possible modification of the NAN model might be making the reversal potential be the variable dependent on intracellular ion concentration, or combine Hodgkin-Huxley model with Goldman-Hodgkin Katz model to describe channel activities. Another limitation of the current NAN models is that we did not explicitly describe the local concentration of Na^+^/Ca^2+^ ions. For example, KCa and KNa channel would sense the ion concentration at the surface of soma, which is not essentially the same as the cytosolic ion concentration.

The NAN model which is described by a set of ordinary differential equations is too complex to be analytically solved. Thus our conclusion derived by the model mostly relied on the numerical computing. Although for detailed analysis we used over 1,000 solutions showing SWS firing pattern in order to guarantee the generalizability of the statements, we cannot exclude the possibility that some of the conclusion shown here is depending on the specific parameters used in the numerical simulation rather than the structure of ODEs.

## Supporting information

Supplementary figures and tables, and will be used for the link to the file on the preprint site.

## ACKNOWLEDGEMENTS

We would like to thank F. Tatsuki for valuable discussion. This work was supported by a Grant-in-Aid for Scientific Research (S) (JSPS KAKENHI, 18H05270, to H.R.U.); a Grant-in-Aid for Transformative Research Areas (A) (JSPS KAKENHI 24H02305, to K.L.O.); a Grant-in-Aid for Scientific Research (C) (JSPS KAKENHI 23K05738, to K.L.O.); Human Frontier Science Program (HFSP) Research Grant Program (RGP0019/2018, to H.R.U.); Exploratory Research for Advanced Technology (ERATO) (JST, JPMJER2001, to H.R.U.); Quantum Leap Flagship Program (Q-LEAP MEXT, JPMXS0120330644, to H.R.U.); Brain Mapping by Integrated Neurotechnologies for Disease Studies (Brain/MINDS) (AMED, JP21dm0207049, to H.R.U.); Innovative Drug Discovery and Development (AMED, JP19am0401011, to H.R.U.); and an intra-mural Grant-in-aid (RIKEN BDR, to H.R.U.).

## AUTHOR CONTRIBUTIONS

T.R.S., K.L.O., and H.R.U. designed the study. T.R.S. performed most of the simulation study. F.L.K. and T.R.S. wrote the code for the network simulation. T.R.S., K.L.O., and H.R.U. analysed the mechanism of SWS oscillation and T.R.S., K.L.O., and H.R.U. wrote the manuscript.

## METHODS

Software used in this study

The model is simulated using Python (version 3.7.0 or version 3.8.5), with odeint function. Subsequent analyses with NAN model and its derivative models were conducted with these following libraries:

matplotlib:3.7.5

numpy: 1.21.6 or 1.23.4

pandas: 0.23.4 or 2.0.3

requests: 2.19.0 or 2.31.0

scikit-learn: 1.3.2

seaborn: 0.13.2

scipy: 1.1.0 or 1.5.2

Network analyses were conducted with GPU in NVIDIA HPC SDK 22.11. Codes were written with C++17 and CUDA 11.8.

## RESOURCE AVAILABILITY

Lead contact

Further information and requests for resources should be directed to and will be fulfilled by Prof. Hiroki R. Ueda with.

### Materials availability

This study did not generate new materials.

### Data and code availability

No raw data was collected for this study. All results were produced from deterministic simulations and can be reproduced by using the same parameters.

All of the code used for simulations and analyses can be found at this link. This URL is going to be replaced upon publication.

For any additional questions or information, please contact the lead contact.

## METHOD DETAILS

### Formulation of several mathematical models for SWS firing pattern with Na^+^ dynamics

Models and parameters used in this paper are based on the AN model^7^. The AN model is constructed based on previous studies ^4-6,10,36,37^ by using Hodgkin-Huxley-type equations.

The NAN model, the NAN model without voltage-gated Ca^2+^ channel, the NAN model with Na^+^/K^+^ ATPase, and the FNAN model are constructed as follows. Formula for the NAN model is given by

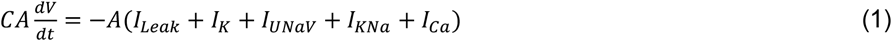

Formula for the NAN model without voltage-gated Ca^2+^ channel is given by

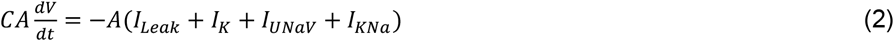

Formula for the NAN model with Na^+^/K^+^ ATPase is given by

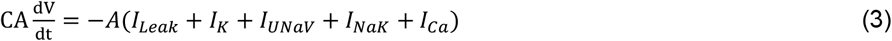

Formula for the FNAN model is given by

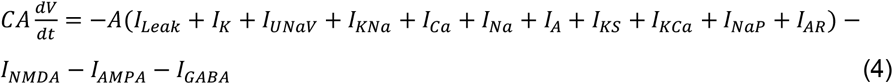

For all the models, C is the membrane capacitance, A is the area of a single neuron, V is the membrane potential, and I is the electrical current of each channel. The other parameters used in this study are listed in **Table S1-5**. Each intrinsic current is given by the following equations (5) to (45):

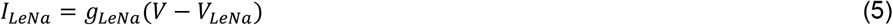

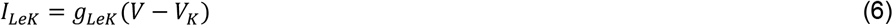

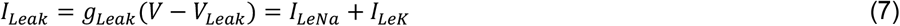

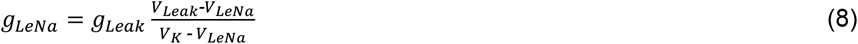

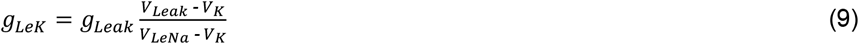

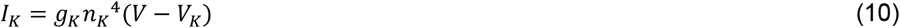

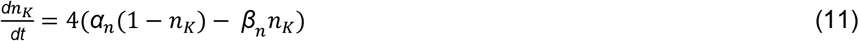

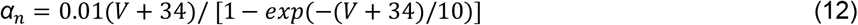

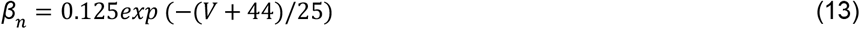

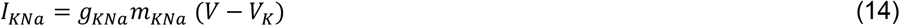

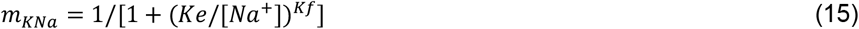

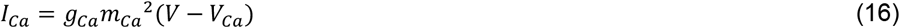

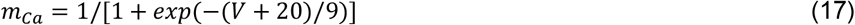

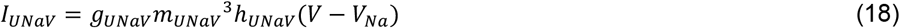

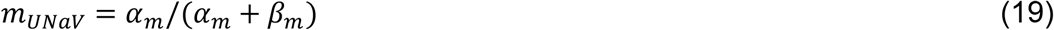

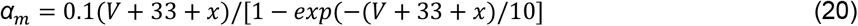

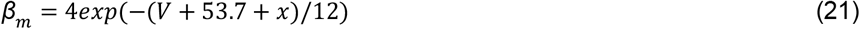

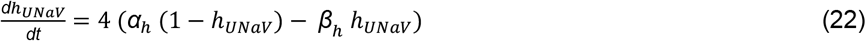

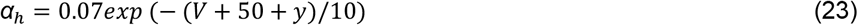

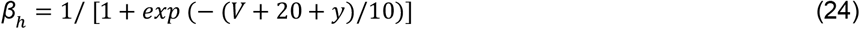

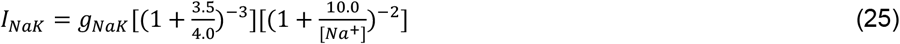

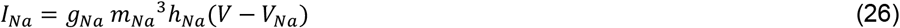

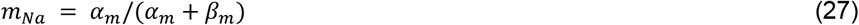

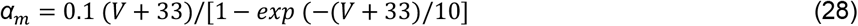

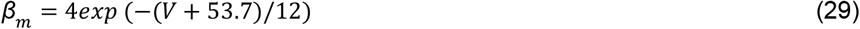

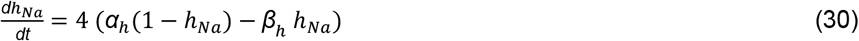

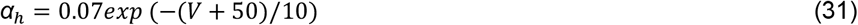

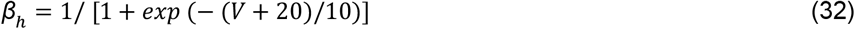

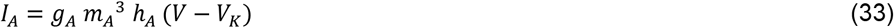

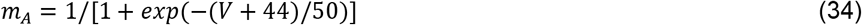

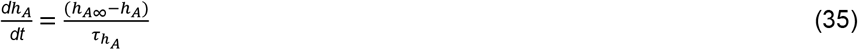

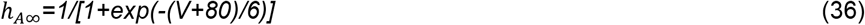

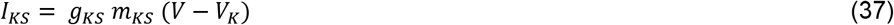

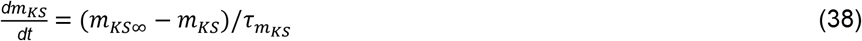

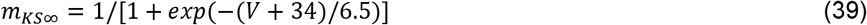

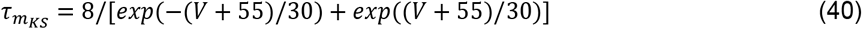

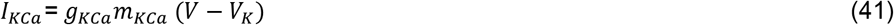

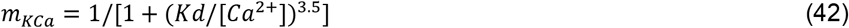

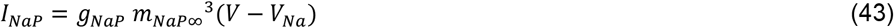

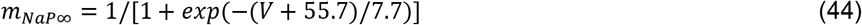

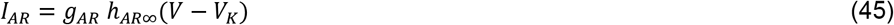

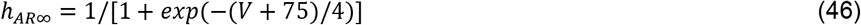

Note that *K*_*e*_ and *K*_*f*_ in the formula of KNa channels (15) is fitted based on the activation curve, **Figure 2b** in the previous study^12^. The formulation of current mediated by Na^+^/K^+^ ATPases (25) is incorporated from the previous study^16^.

In the FNAN model, the description of synaptic elements is altered from the AN model so that they are only influencing the membrane potential and they don’t influence intracellular ion concentration. This assumption is different from the assumption in the AN model that synaptic currents alter the intracellular ionic concentration^7^. This is because previous study suggested that the ionic diffusion across the spine is negligible^38^, and the spine head functions as a separate compartment from the soma. Therefore, in FNAN model, the action of NMDAR and AMPAR are described to be unrelated to intracellular Na^+^ dynamics, and NMDAR is described to be unrelated to intracellular Ca^2+^ dynamics. Each extrinsic current is then given by the following equations (47) to (56):

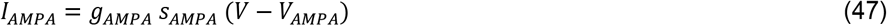

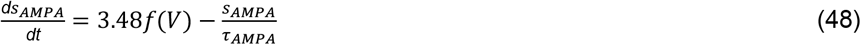

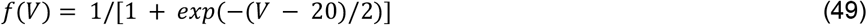

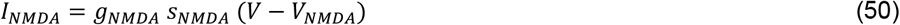

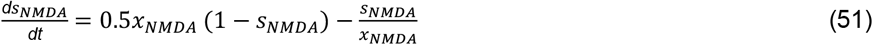

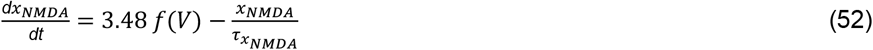

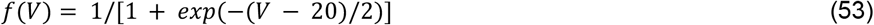

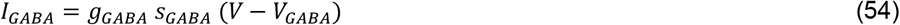

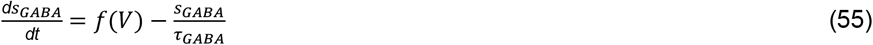

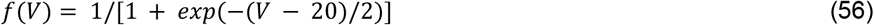

Leak Na channels (NALCN channels) allow Na^+^, K^+^, Ca^2+^ currents^39^. Therefore, leak Na^+^ current can be divided into Na^+^, K^+^, Ca^2+^ currents as follows:

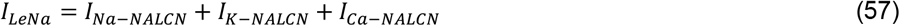

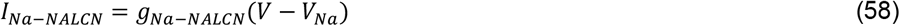

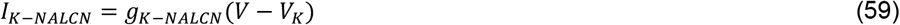

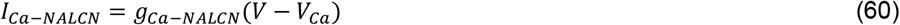

These must satisfy the following equation.

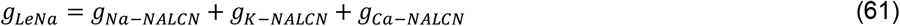

*g*_*LeNa*_ = *g*_*Na*−*NALCN*_ + *g*_*K*−*NALCN*_ + *g*_*Ca*−*NALCN*_ (61) In order to determine the proportion of *g*_*Na*−*NALCN*_, *g*_*K*−*NALCN*_, and *g*_*Ca*−*NALCN*_, electrophysiological recordings of NALCN channels are used^39^. Suppose the reversal potential of NALCN channels is *γ*_1_ mV when [*Na*^+^]_*o*_ = 155 mM^39^. Suppose the reversal potential of NALCN channels is *γ*_2_ mV when [*Na*^+^]_*o*_ = 15.5 mM^39^. Assuming that the reversal potential of Na^+^ is *V*_*Na*1_ when [*Na*^+^]_*o*_ = 155 mM, and the reversal potential of Na^+^ is *V*_*Na*2_ when [*Na*^+^]_*o*_ = 15.5 mM, based on the definition of reversal potential, the following equations hold.

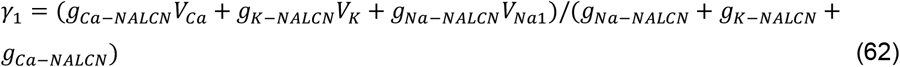

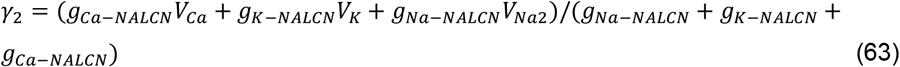

Therefore,

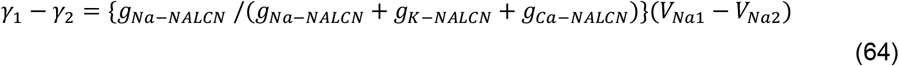

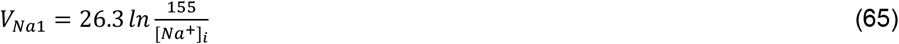

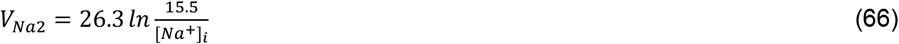

Therefore,

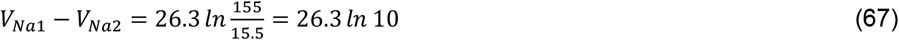

Therefore,

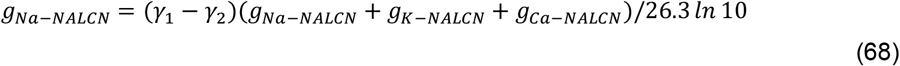

Suppose the reversal potential of NALCN channels is *δ*_1_mV when [*K*^+^]_*o*_ = 155 mM^39^. Suppose the reversal potential of NALCN channels is *δ*_2_ mV when [*K*^+^]_*o*_ = 15.5 mM^39^. Assuming that the reversal potential of K^+^ is *V*_*K*1_ when [*K*^+^]_*o*_ = 155 mM, and the reversal potential of K^+^ is *V*_*K*2_ when [*K*^+^]_*o*_ = 15.5 mM, based on the definition of reversal potential, the following equations hold.

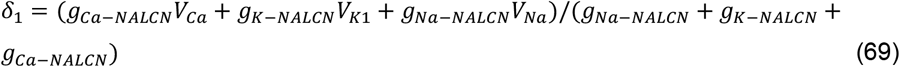

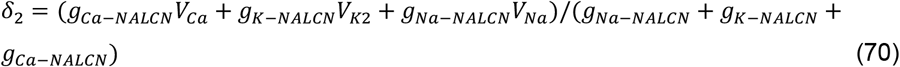

Therefore,

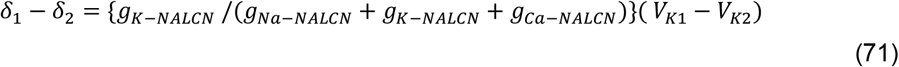

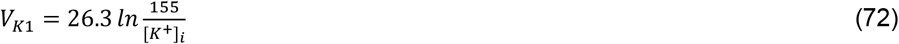

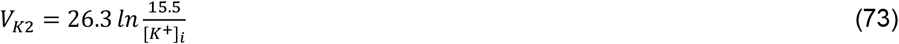

Therefore,

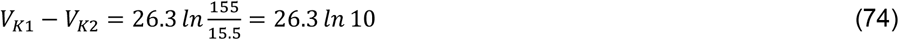

Therefore,

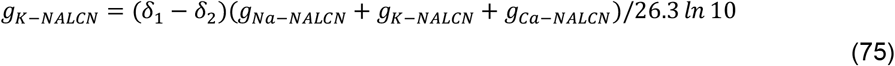

Suppose the reversal potential of NALCN channels is ϵ_1_ mV when [*Ca*^2+^]_*o*_ = 100 mM^39^. Suppose the reversal potential of NALCN channels is ϵ_2_ mV when [*Ca*^2+^]_*o*_ = 10 mM^39^. Assuming that the reversal potential of Ca^2+^ is *V*_*Ca*1_ when [*Ca*^2+^]_*o*_ = 100 mM, and the reversal potential of Ca^2+^ is *V*_*Ca*2_ when [*Ca*^2+^]_*o*_ = 10 mM, based on the definition of reversal potential, the following equations hold.

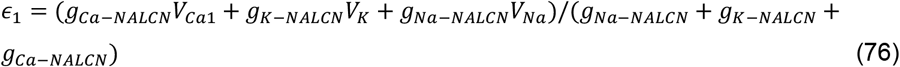

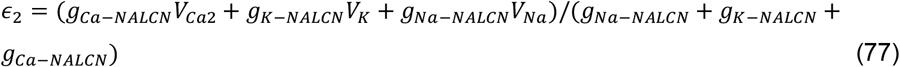

Therefore,

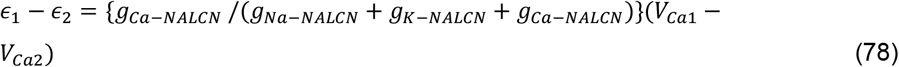

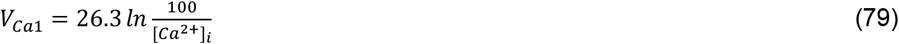

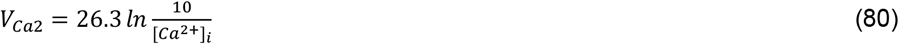

Therefore,

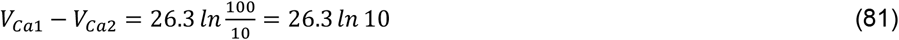

Therefore,

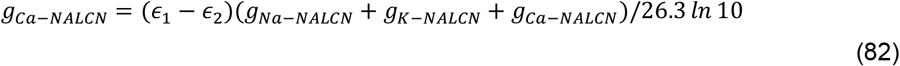

Based on the equation (68), (75) and (82), the proportion of g_Na-NALCN_, g_K-NALCN_, and g_Ca-NALCN_ is determined as follows:

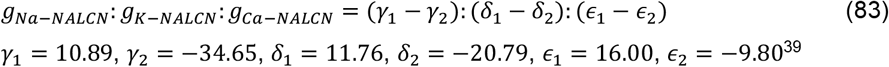

Therefore,

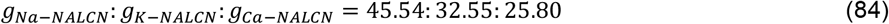

Therefore,

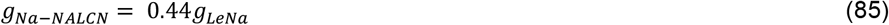

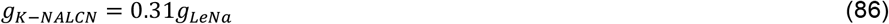

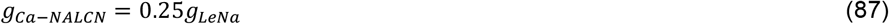

The time course of intracellular Na_+_ concentration is also modeled as follows: Formula for the NAN model and the NAN model without voltage-gated Ca^2+^ channel are given by

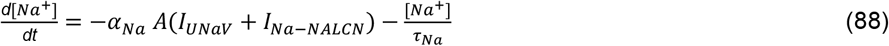

Formula for the NAN model and the NAN model with Na^+^/K^+^ ATPase isgiven by

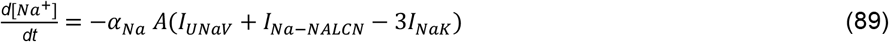

Formula for FNAN model is given by

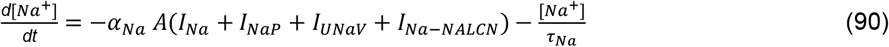

The time course of intracellular Ca^2+^ concentration in the FNAN model was also modeled as follows:

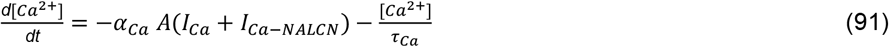

The coefficient of Na^+^-entry (αNa) is estimated under the assumption that the average volume of a single neuron is ∼10 pl^40^. The increase in intracellular Na^+^ concentration caused by Na^+^ currents was determined as follows: 1 nA Na^+^ current for 1 ms causes 1 nA×1 ms/ ∼10 pl =1 pC/ ∼10 pl = ∼1.0 /10 F mol/L = ∼ 1.0 μM increase in the intracellular Na^+^ concentration, where F = ∼0.96 × 10^5^ C/mol is the Faraday constant.

Integration is performed with the following initial values in each parameter search, and bifurcation analysis: V = -45 mV, h_NaV_ = 0.045 (unitless), h_UNaV_=0.045 (unitless), n_K_ = 0.54 (unitless), [Na^+^] = 1 mM.

### Parameter search

We searched for parameter sets with SWS firing patterns. Parameter sets are randomly created in a parameter space defined below: the conductance of intrinsic (non-synaptic) channels in the soma (*g*_Leak_, *g*_Na_, *g*_KNa_, *g*_UNaV_, *g*_K_, *g*_NaK_, *g*_Ca_, *g*_A_, *g*_KS_, *g*_KCa_, *g*_NaP_, *g*_AR_) and extrinsic (synaptic) channels in the dendrite (*g*_AMPA_, *g*_NMDA_, *g*_GABA_) are generated by selecting parameters from an exponential distribution bounded to the interval 0.001-10 mS/cm^2^, time constant of Na^+^ pumps/exchangers (*τ*_Na_) is generated by selecting a parameter from an exponential distribution bounded to the interval 1000-10000 ms, time constant of Ca^2+^ pumps/exchangers (*τ*_Ca_) is generated by selecting a parameter from an exponential distribution bounded to the interval 10-1000 ms, shift of the activation curve (*x*) and the inactivation curve (*y*) are generated by selecting parameters from a uniform distribution bounded to the interval -45-45 mV. The intervals used in the parameter searches for the conductance of the intrinsic and extrinsic channels are set to be the same range after the consideration of the area of a neuron, A = 0.02 mm^2^ (i.e. 0.0002-2 μS/ 0.002 mm^2^ = 0.001-10 mS/cm^2^). The procedure of parameter search is the same as the previous study^7^ except for *g*_UNaV_, *g*_KNa_, *g*_NaK_, *τ*_Na_, *x*, and *y*, which are newly introduced in this paper. Differential equations are solved to compute the membrane potential from 0 to 20 seconds. The simulated time course of membrane potential from 10 to 20 seconds are specifically used for analyzing the wave pattern of membrane potential. This interval is chosen to exclude the influence of the initial conditions of the differential equations on the observed wave pattern. The procedure of wave-type classification is the same as the previous study^7^. The major frequency of the oscillatory behavior is then analyzed by Fast Fourier transform. We also evaluated the fine structure of the wave pattern by counting the number of spikes per 1 second; the number of spikes is determined as half the number of times the membrane potential crossed -20 mV. Solutions, in which the membrane potential exceeded this threshold at almost all time points (> 95%), are eliminated at this point (labeled as “ELSE”). Based on these characteristics, the solutions are classified into four categories: “RESTING” (spike numbers per second < 2 or peak frequency = 0 Hz), “SWS” (0 Hz < peak frequency < 10 Hz and spike number per second > 5 × peak frequency), “AWAKE” (peak frequency ≥ 10 Hz), and “SWS with few spikes” (0 Hz peak frequency <10 Hz and spike number per second < 5 × peak frequency). “SWS with few spikes” indicated that the solution represented slow-wave activity, with fewer than five spikes during one bursting phase of neural activity. All of the solutions classified as “SWS” are then checked manually to select the ones that exhibit oscillatory membrane potential alternating between bursting and silent phases.

### Bifurcation analysis

To analyze the behavior of the system around the parameter set that gave SWS firing patterns, we conducted bifurcation analyses. Each conductance or the time constant was gradually changed from 0.001 to 10 times its original value; note that for *x*, and *y*, the range of the gradual changes are from -45 to +45 of the original value. The solutions were then classified into four categories (RESTING, SWS, AWAKE, ELSE) as described above. In the bifurcation analysis, “SWS” (SWS) components include both “SWS” and “SWS with few spikes”. Manual curation of the classification was not conducted for the result of bifurcation analysis.

### Plotting the normalized Voltage (V) and [Na^+^]

In order to see the link between V and [Na^+^] time courses, we plotted the time-course for all parameter sets yielding SWS firing pattern. For V, the maximum voltage is expressed by 3 and minimum voltage was expressed by -3. For [Na^+^], the maximum concentration is expressed by 3 and minimum concentration is expressed by -3. The period of each oscillation is standardized for all parameter sets and plotted.

### Choosing the representative parameter set

Principal Component Analysis (PCA) is conducted to the parameter with ∼1,000 SWS firing pattern. Subsequently, the probability density function is calculated for PC1 and PC2 based on Kernel Density Estimation by using PCA function in scikit-learn library. Five parameter sets were chosen according to the probability density (top five), and the representative parameter set was chosen arbitrary from the five parameter sets.

### Plotting the intersection of nullplanes

NAN model is composed from ODEs with 4 variables (V, n_K_, h_UNaV_, Na^+^). By fixing the value of the intracellular Na^+^, it can be converted to ODEs with 3 variables (V, n_K_, h_UNaV_). By using 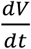 described in (1) and 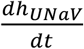 described in (22), intersection of V, h_UNaV_ nullplanes satisfies 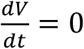 and 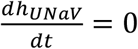, given by the equation:

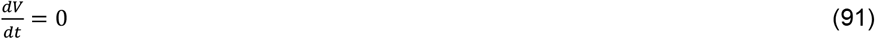

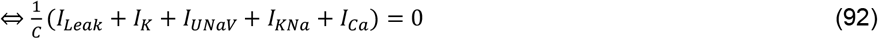

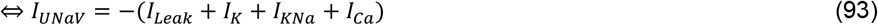

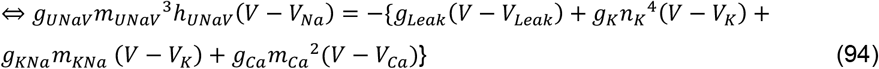

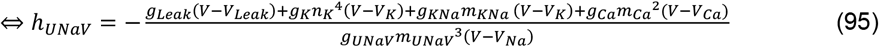

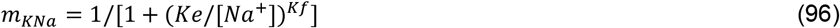

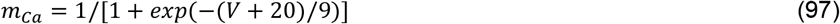

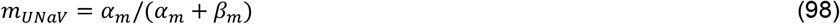

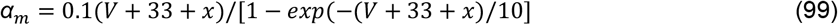

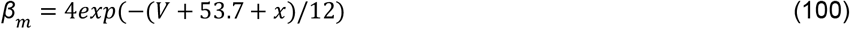

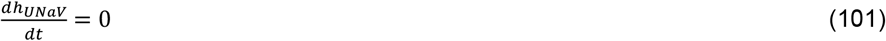

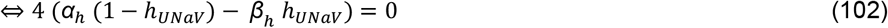

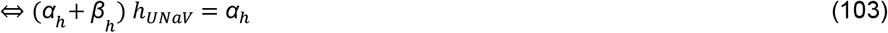

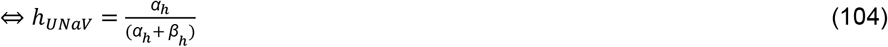

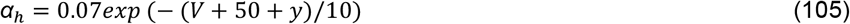

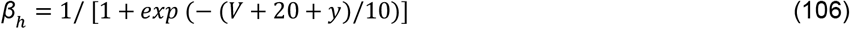

Therefore, if the value of the membrane voltage (V) is determined, the value of n_K_ and h_UNaV_ are determined as follows.

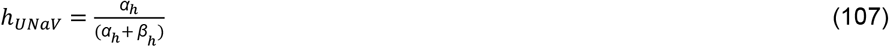

n_K_ satisfies the following equation.

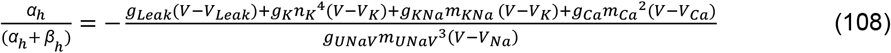

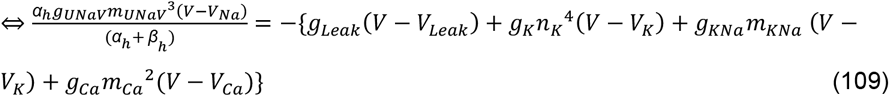

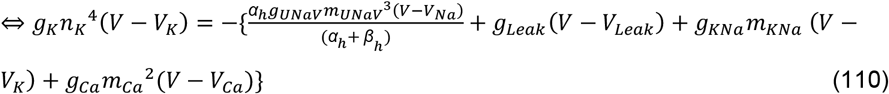

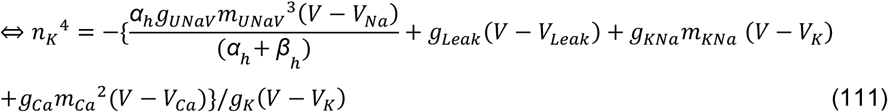

Therefore,

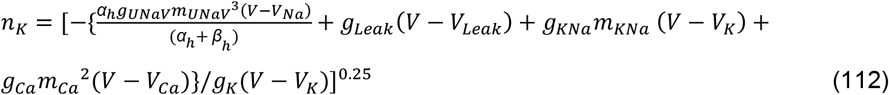

By using 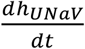 described in (22) and 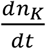 described in (11), intersection of n_K_, h_UNaV_ nullplanes are the line that satisfies 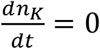 and 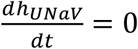.

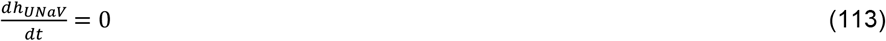

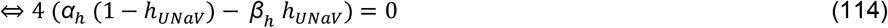

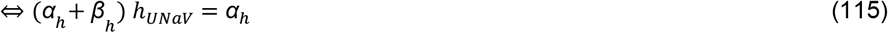

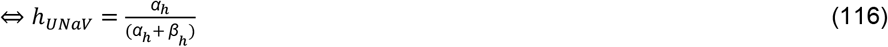

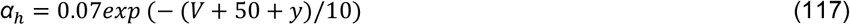

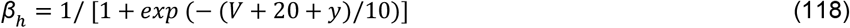

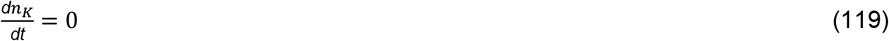

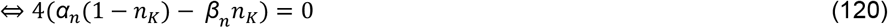

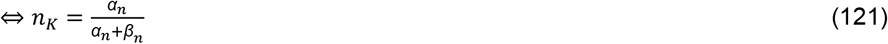

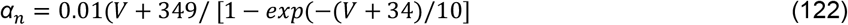

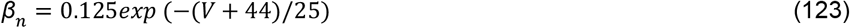

Therefore, if the value of the membrane voltage (V) is determined, the value of n_K_ and h_UNaV_ are determined as follows.

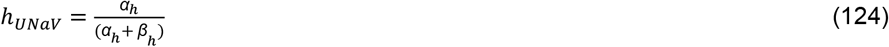

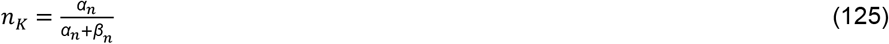

### Measuring up state duration, down state duration, Na^+^ oscillation amplitude, and ISI

The duration of the up state and the down state is determined as follows (**Figure S3A**). In a two-dimensional plane where the x-axis represents time (t) and the y-axis denotes the membrane potential (V), V = f(t) The time course of the membrane potential is given by V = f(t).

First, using Fast-Fourier transform (FFT) analysis on the V = f(t) time series data, the frequency of up-down oscillation is determined as the frequency *fq* (Hz) with the highest power (**Figure S3A**).

Then, the number of intersection points between V = f(t) and V = k (k is increased from -110mV to 40mV in increments of 0.1mV) was computed. When the value of k is small, the intersection points would lie in the down state (**Figure S3B**). By gradually increasing the value of k, the intersection points shift toward the up state, where the number of intersection points shall be increased abruptly (**Figure S3C**). Hence, by examining the relationship between the value of k and the number of intersection points, the membrane potential, at which the transition between the up state and the down state occurs can be estimated as the k value with acute increase of the intersection points. The detailed algorithm was implemented as follows.

We define the threshold value of k = k_thresh_, as the up state membrane voltage close to the boundary between up/down state transition. k_thresh_ was calculated as follows: as the value of k is increased from -110 mV in increments of 0.1 mV, the number of intersection point gradually increases. Then, right after the number of intersection point exceeds 30 × 2 × *fq* + 100, the value of k is determined as k_thresh_. This is because 30 × 2 × *fq* matches with the number of up/down state transition timing during the simulated time of 30 sec and addition of 100 makes sure the marked increase of intersection points within the up state.

Next, among all the intersection points between V = f(t) and V = k_thresh_, intersection points accounting for the transition from the down state to the up state or from the up state to the down state are selected. t coordinate of intersection points are defined as t_int_ (msec), and are determined as follows:

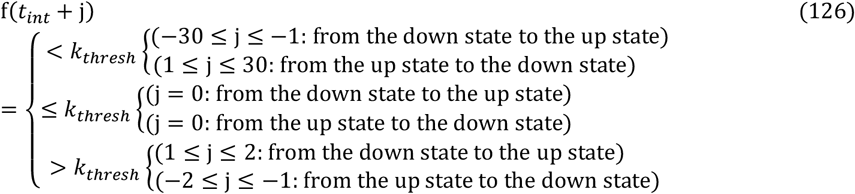

Transition point between the up state and the down state are determined as above because at the transition point from the down state to the up state, f(t) is constantly below k_thresh_ when t_int_<t and only after t exceeds t_int_, f(t) is larger than k_thresh_ because during the down state membrane potential is hyperpolarized and only when it is in the up state period membrane potential surpasses the threshold voltage (**Figure S3D**). The same logic is applied when determining the transition point from the up state to the down state (**Figure S3E**).

By using the transition time between the up state and the down state, up state period and down state period are calculated. There periods are calculated for many cycles of oscillation, so the average period of the up state and the down state are calculated.

Na^+^ oscillation amplitude is calculated for many cycles of oscillation, so the average Na^+^ oscillation amplitude is calculated.

To calculate ISI, firstly all local maximum points are extracted from the time-series data of V=f(t) by using argrelmax function in SciPy library. ISI is defined as the interval between adjacent local maximum points. However, ISI exceeding 60 msec is excluded because too long ISI can be confused with the short down state. ISI accounting for all adjacent spikes are calculated individually, and the average ISI is calculated.

### Construction of the network model

A Network model consisted of 84 neurons is constructed. The parameters (i.e. conductance of ion channels and synaptic receptors) are homogeneous throughout all neurons in one network model. Each neuron is randomly connected to other neurons with a probability of 20% by AMPA-and NMDA-mediated synapses. Out of all 2,137 parameter sets with SWS firing pattern in the FNAN model, 1703 parameter sets showed SWS firing pattern in the FNAN model where *g*_GABA_ = 0. Among them, we chose 1,057 parameter sets with SWS firing pattern in the FNAN model which are confirmed by calculation with a forth-order Runge-Kutta method with a time step of 0.1 msec. Finally, 100 parameter sets are chosen as representative parameter sets out of these 1,057 parameter sets. The way representative parameter sets are selected is the same as the way representative parameter sets are selected in the analysis using an averaged neuron with mean-field approximation.

Simulations were conducted using GPU (NVIDIA GeForce RTX 3090) and numerical calculations were conducted by a forth-order Runge-Kutta method with a time step of 0.01 msec.

