## Supplementary figures and tables, and will be used for the link to the file on the preprint site. for "A Design Principle for slow-wave sleep firing pattern with Na^+^ dynamics"

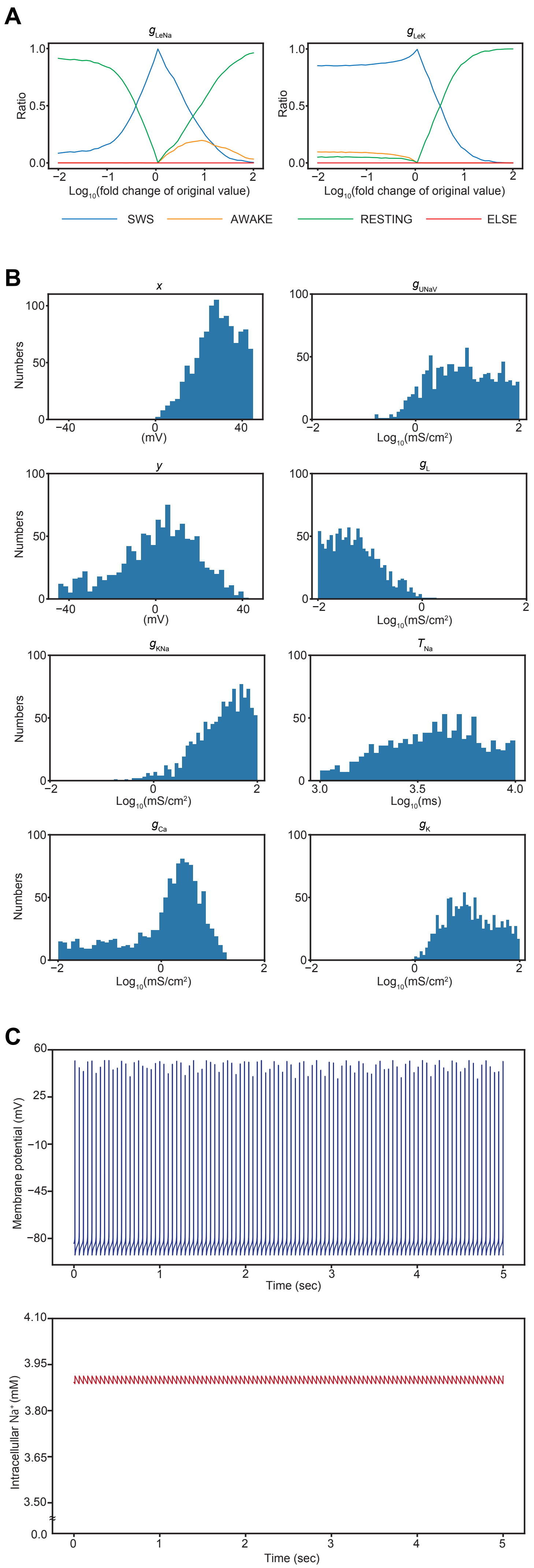


Supplementary Figure 1. Construction of the NAN model by incorporating UNaV channel.

#### Supplementary Figure 1. Construction of the NAN model by incorporating UNaV channel.

(A) The results of the bifurcation analysis with $g_{LeK}$, $g_{LeNa}$ in the NAN model.

(B) The distribution of parameters with SWS firing pattern.

(C) A example case of AWAKE firing patterns was generated by upregulating $g_{K}$ in **Figure 1E**.

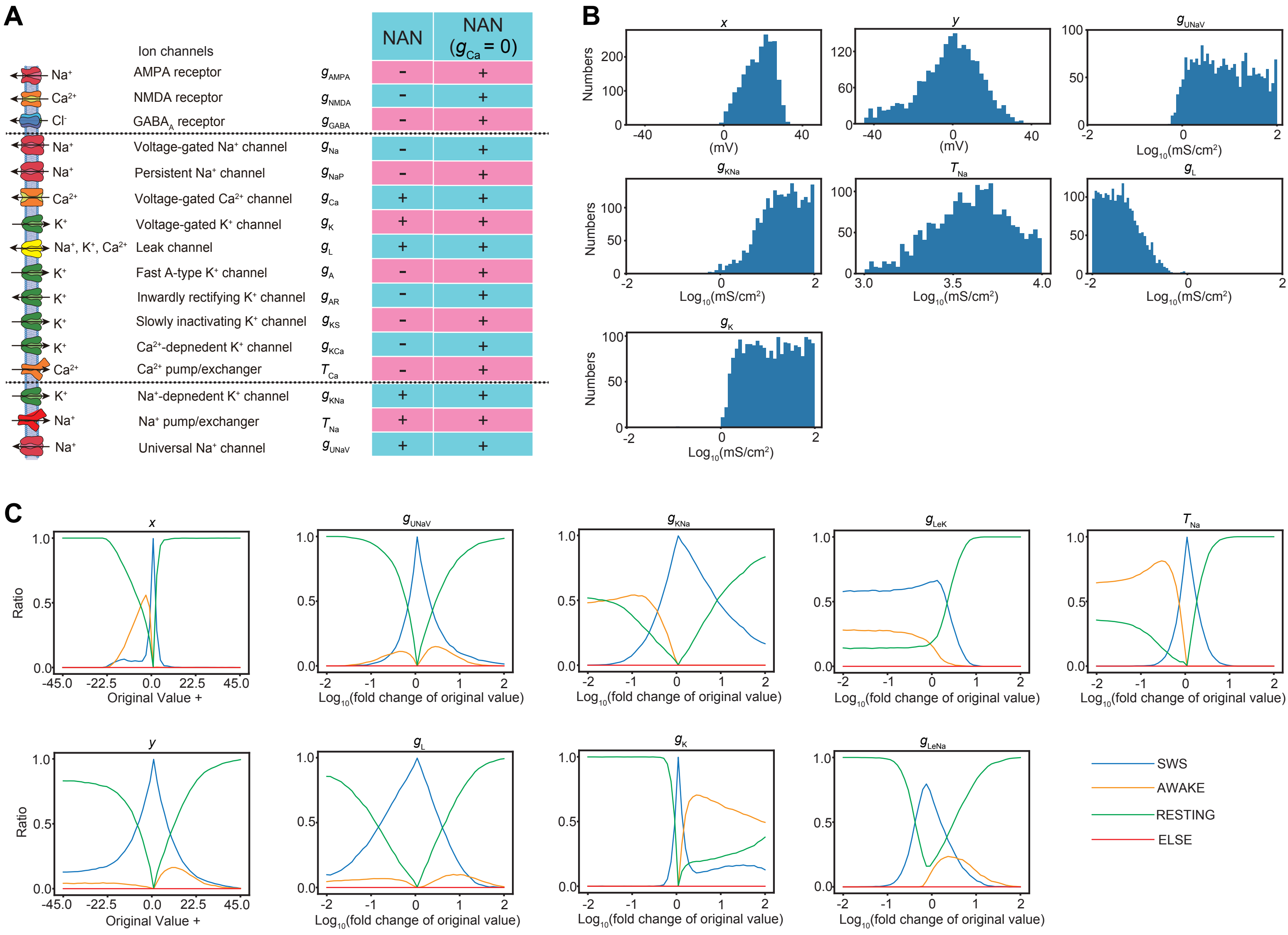


Supplementary Figure 2. Construction of the NAN without voltage-gated $Ca^{2+}$ channel.

#### Supplementary Figure 2. Construction of the NAN model without voltage-gated $Ca^{2+}$ channel.

(A) Schematic diagram of the NAN model and the NAN model without voltage-gated $Ca^{2+}$ channels (labeled as the NAN model with $g_{Ca} = 0$).

(B) The distribution of parameters with SWS firing pattern in the NAN model without voltage-gated $Ca^{2+}$ channels.

(C) The results of the bifurcation analysis in the NAN model without voltage-gated $Ca^{2+}$ channels.

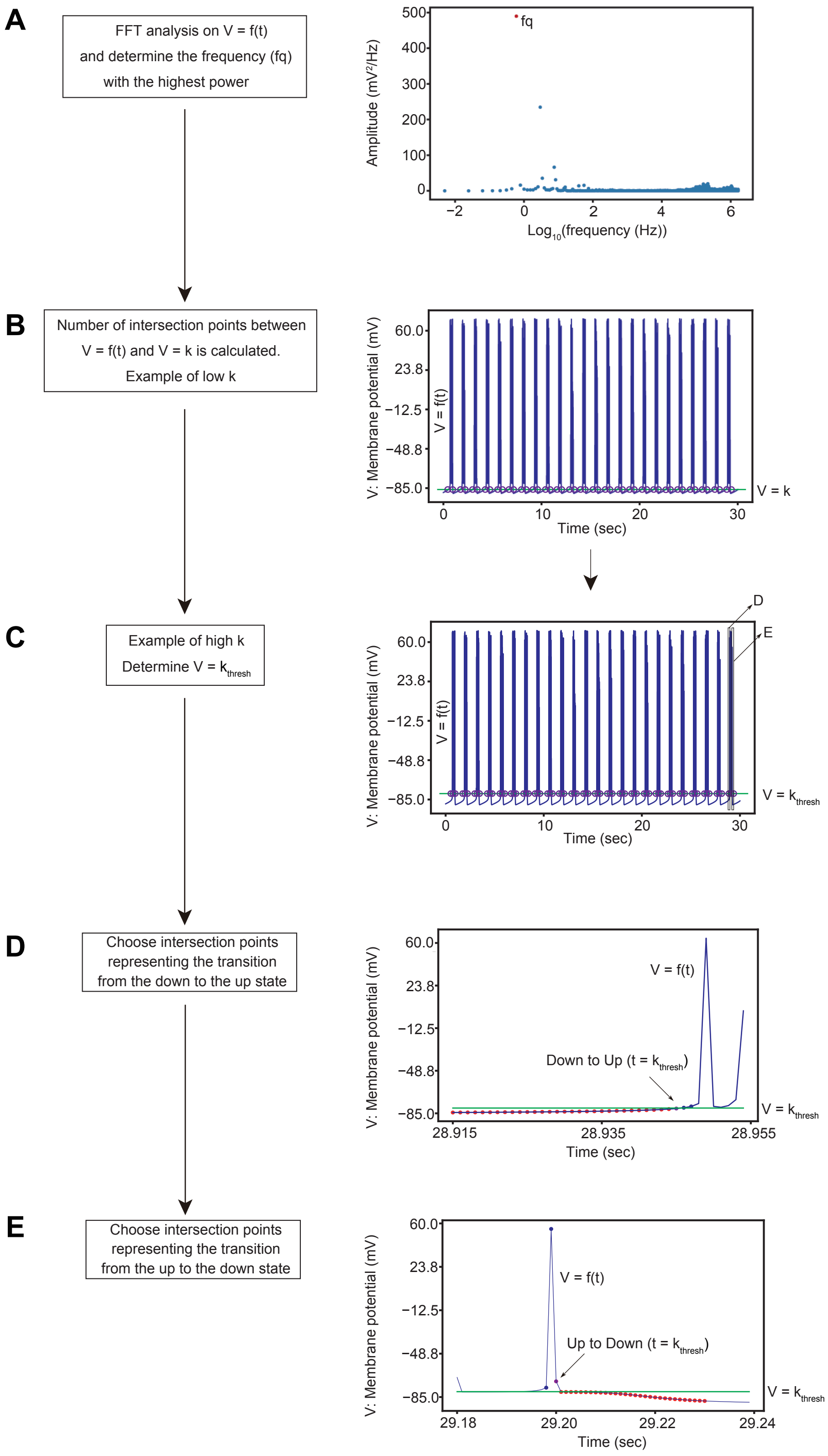


Supplementary Figure 3. Activation parameter of UNaV is important for controlling several features of wave pattern in the NAN model.

**Supplementary Figure 3. Activation parameter of UNaV is important for controlling several features of wave pattern in the NAN model.**

(A) Power spectrum diagram. Frequency with the highest power is selected.

(B) The relation between V=k (green) and V=f(t) (blue). The intersection points encircled by the purple circles are in the down state.

(C) The relation between $V=k_{thresh}$ (green) and V=f(t) (blue). The intersection points encircled by the purple circles are in the up state. $V = k_{thresh}$ is at the bottom of the up state, so the intersection points include transition points between the up state and the down state.

(D) Amongst all purple points in (C), points which denote the transition from the down state to the up state are selected. The purple point is amongst all points in (C), and it is just below the line: $V = k_{thresh}$. The blue points have to be above the line: $V = k_{thresh}$ and the red points have to be below the line if the purple point is indeed the transition point from the down state to the up state.

(E) Amongst all purple points in (C), points which denote the transition from the up state to the down state are selected. The purple point is amongst all points in (C), and it is just above the line: $V = k_{thresh}$. The blue points have to be above the line: $V = k_{thresh}$ and the red points have to be below the line if the purple point is indeed the transition point from the up state from the down state.

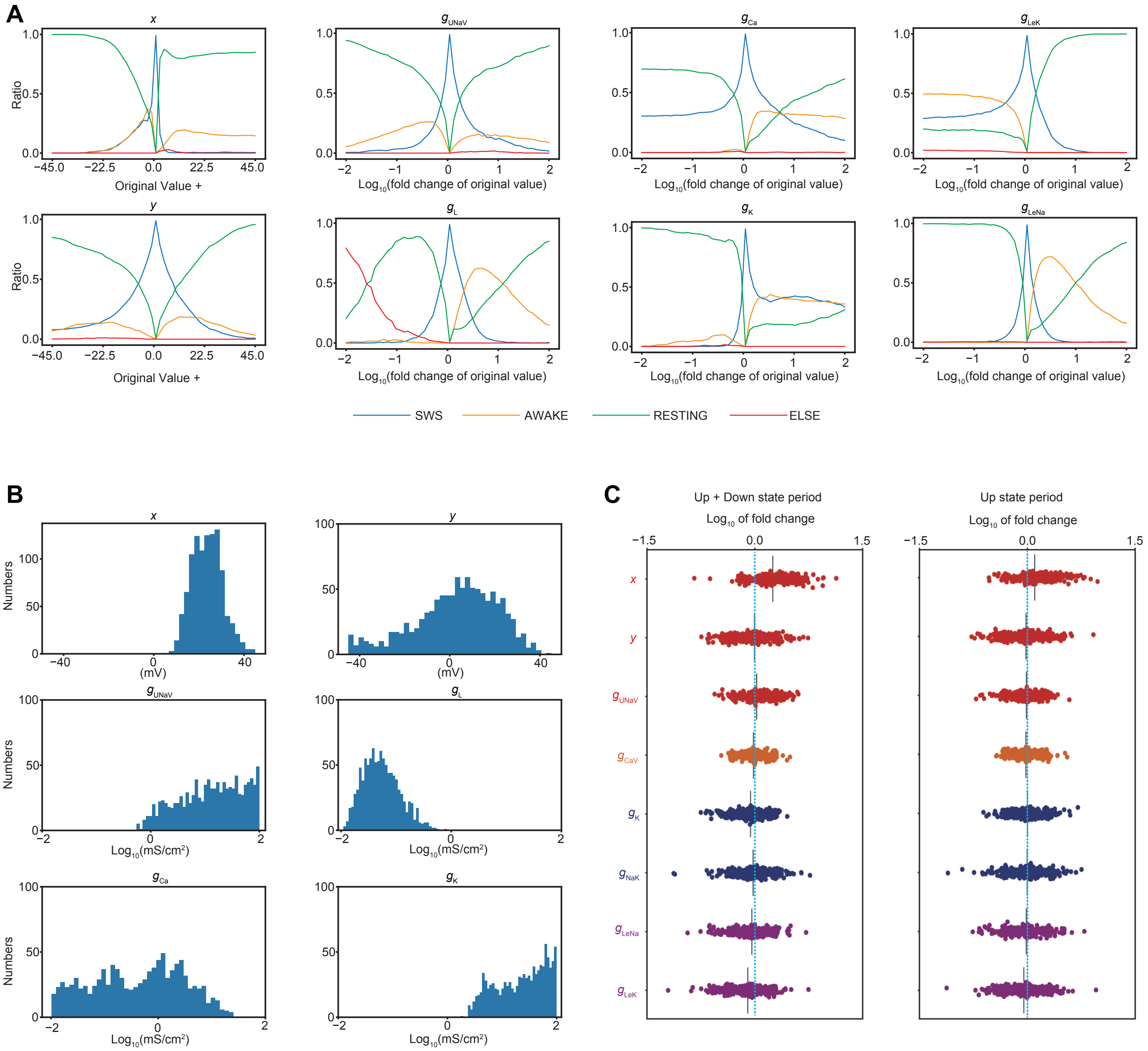


Supplementary Figure 4. $Na^+$-dependent hyperpolarization pathway can also be implemented by $Na^+/K^+$ ATPase.

#### Supplementary Figure 4. $Na^+$-dependent hyperpolarization pathway can also be implemented by $Na^+/K^+$ ATPase.

(A) The results of the bifurcation analysis in the revised model with $Na^+/K^+$ ATPases.

(B) The distribution of parameters with SWS firing pattern in the NAN model with $Na^+/K^+$ ATPases.

(C) The effect of slight upregulation of the value of each parameter alter the characteristics of SWS firing pattern (Up + Down state period, $Na^+$ oscillation amplitude).

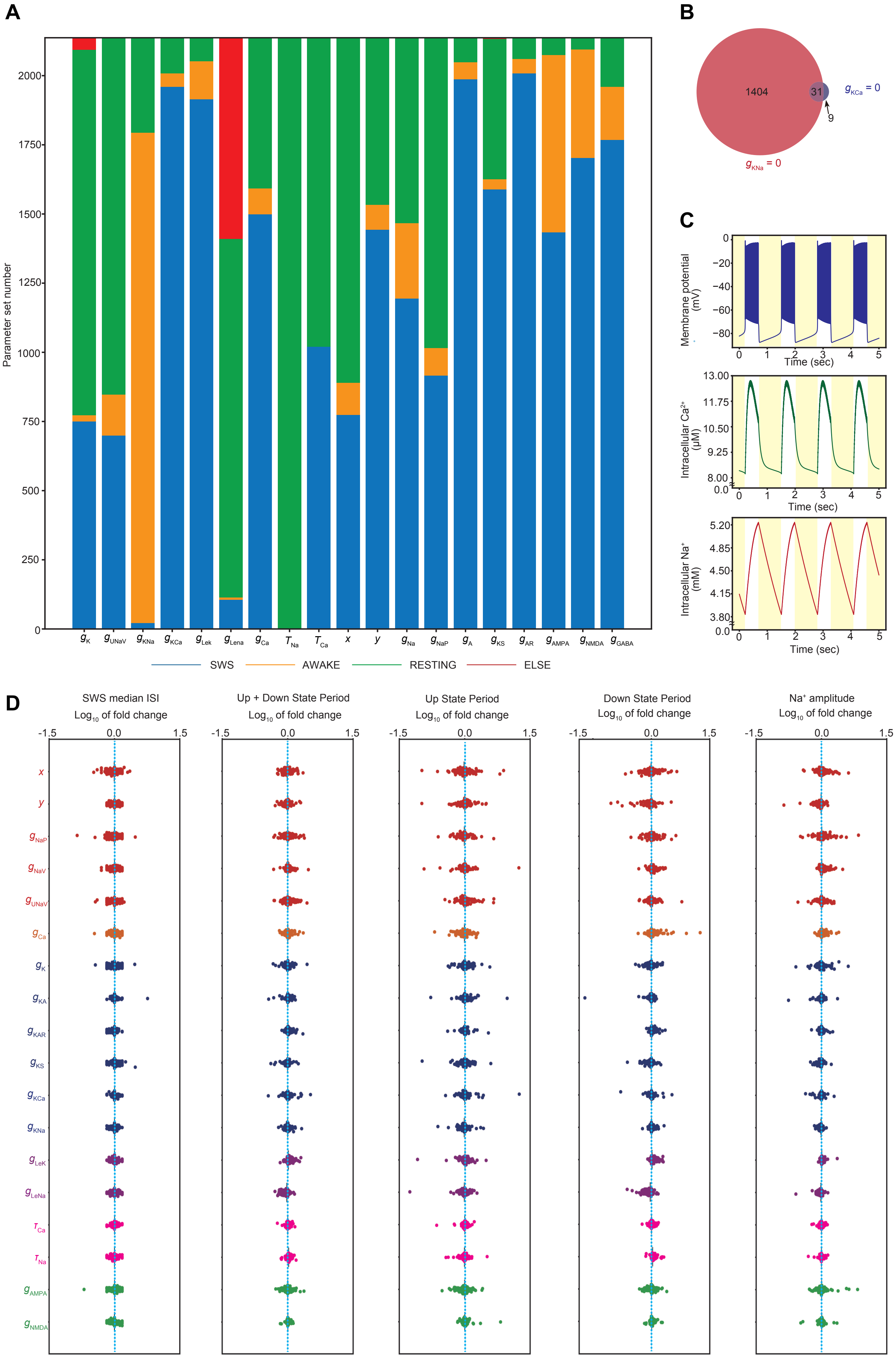


Supplementary Figure 5. The Full-NAN model reveals the importance of $Na^+$ dependent pathway.

#### Supplementary Figure 5. The Full-NAN model reveals the importance of $Na^+$-dependent hyperpolarization pathway.

(A) The result of knockout analysis with all parameter sets showing SWS firing pattern. When the contribution of a particular element is set to zero, the amount of parameter sets showing AWAKE firing pattern is shown by the color, orange, and the amount of parameter sets showing RESTING firing pattern is shown by the color, green and the amount of parameter sets which are excluded from the analysis (ELSE) because of their aberrant firing pattern is shown by the color, red, and the amount of parameter sets showing SWS firing pattern is shown by the color, blue. Classification criteria of SWS, AWAKE, RESTING, ELSE follows that of Tatsuki et al., 2016[7].

(B) A Benn diagram showing how many parameters show AWAKE firing pattern when KNa channels' current or KCa channels' current are downregulated.

(C) Representative SWS firing patterns, intracellular $Ca^{2+}$ concentration, and intracellular $Na^+$ concentration of the FNAN model. Yellow areas denote the down state whereas white areas denote the up state.

(D) The effect of slight upregulation of the value of each parameter alters the characteristics of SWS firing pattern. All parameter sets yielding SWS firing pattern are used in the analysis. For parameter *x* and *y*, characteristics of SWS firing pattern is calculated when the value of *x*, *y* is (i): 0.5 lower than the original value (ii): 0.5 higher than the original value. For the other parameters, characteristics of SWS firing pattern is calculated when the value is (i): 0.975 times of the original value (ii): 1.025 times of the original value. The value calculated in (ii) is divided by the value calculated in (i). Blue dotted line shows $Log_{10}$ of fold change = 0 (i.e. perturbation does not alter each characteristic of SWS firing pattern).

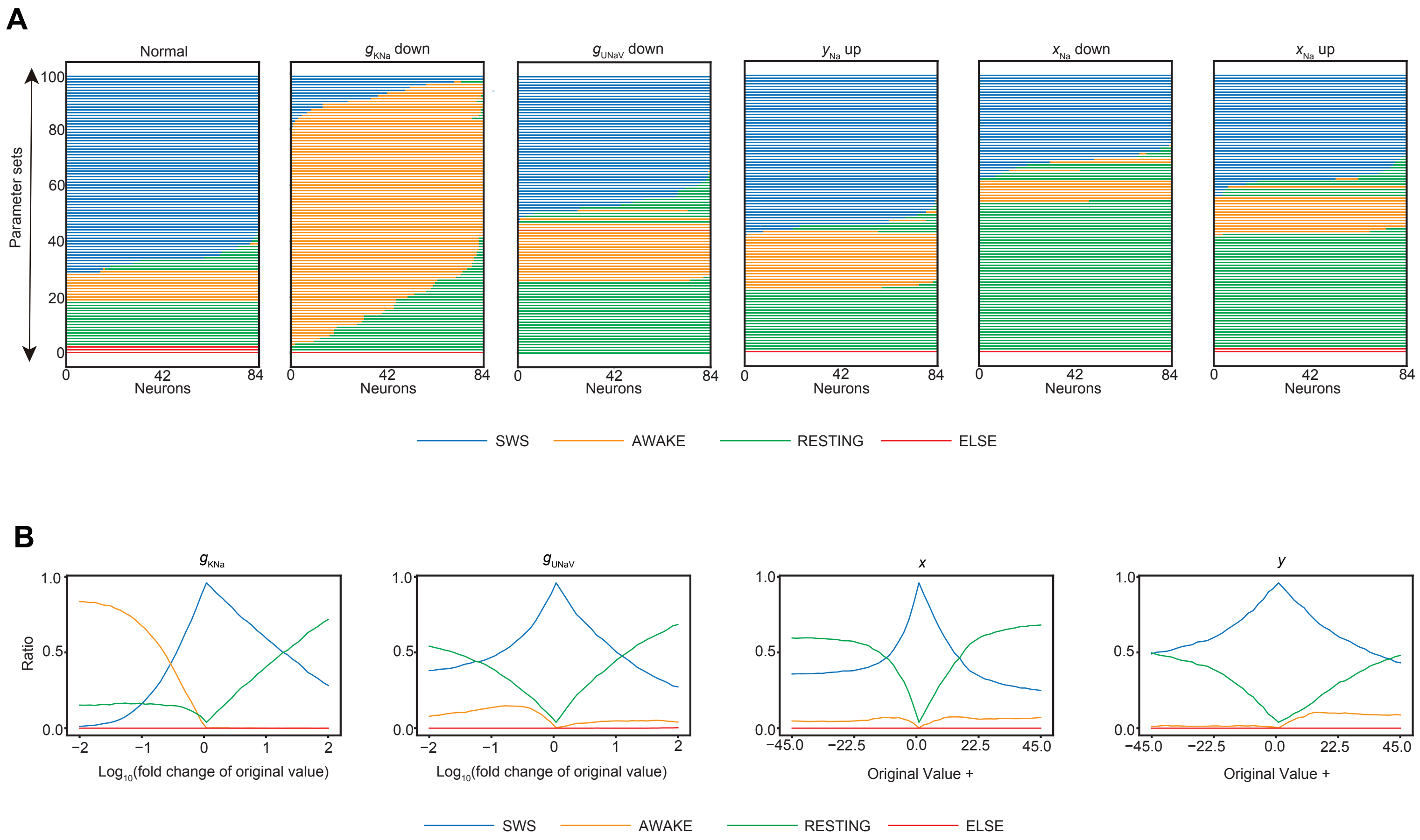


Supplementary Figure 6. The Full-NAN model reveals the importance of $Na^+$ dependent pathway.

#### Supplementary Figure 6. The Full-NAN model reveals the importance of $Na^+$-dependent hyperpolarization pathway.

(A) Changes in the firing pattern of each cell when the value of one parameter indicated above the figure is altered. Perturbation is given to each parameter set, and firing patterns of each neuron in the network are visualized. The exact amount of perturbation given to each parameter is determined by the results of the bifurcation analysis in **Figure S5A** (*x* down; -12.6, *x* up; +12.6, *y*; +14.4, $g_{UNaV}$; $\times 10^{-0.72}$, $g_{KNa}$; $\times 10^{-2.0}$).

(B) The results of the bifurcation analysis in the FNAN model.

### Supplemental tables

Table S1-S5 show parameter values of representative parameter sets in the models used this study.

Table S1. Parameters used in Representative parameter sets in the NAN model.

| Parameter | Description | Value |
|---|---|---|
| C | Membrane Capacitance | 1 μF/cm$^2$ |
| A | Area of Neuron | 0.02 mm$^2$ |
| $V_{Na}$ | Sodium reversal potential | 55 mV |
| $V_K$ | Potassium reverse potential | -100 mV |
| $V_{Ca}$ | Calcium reversal potential | +120mV |
| Ke | Dissociation constant of KNa channels | 32 μM |
| Kf | Parameter in KNa channels | 3 |
| $V_{LeNa}$ | Leak $Na^+$ reversal potential | 0 mV |
| $\alpha_{Na}$ | Coefficient of $Na^+$ entry | 1.0 μM/(nA ms) |
| $g_K$ | The conductance of the voltage-gated $K^+$ channels | 9.956981784 mS/cm$^2$ |
| $g_{UNaV}$ | The conductance of the UNaV channels | 11.47684485 mS/cm$^2$ |
| $g_{KNa}$ | The conductance of KNa channels | 29.01862745 mS/cm$^2$ |
| $g_{Leak}$ | The conductance of the leak channels | 0.030405956 mS/cm$^2$ |
| $g_{Ca}$ | The conductance of the voltage-gated $Ca^{2+}$ channels | 4.79650149 mS/cm$^2$ |
| $\tau_{Na}$ | The time constant of $Na^+$ efflux | 5352.42370631 ms |
| $x$ | Shift of the activation curve | 28.96364873 mV |
| $y$ | Shift of the inactivation curve | 1.11129524 mV |

Table S2. Parameters used in Representative parameter sets in the NAN model with $Na^+/K^+$ ATPase.

| Parameter | Description | Value |
|---|---|---|
| C | Membrane Capacitance | 1 μF/cm$^2$ |
| A | Area of Neuron | 0.02 mm$^2$ |
| $V_{Na}$ | Sodium reversal potential | 55 mV |
| $V_K$ | Potassium reverse potential | -100 mV |
| $V_{Ca}$ | Calcium reversal potential | +120 mV |
| Ke | Dissociation constant of KNa channels | 32 μM |
| Kf | Parameter in KNa channels | 3 |
| $V_{LeNa}$ | Leak $Na^+$ reversal potential | 0 mV |
| $\alpha_{Na}$ | Coefficient of $Na^+$ entry | 1.0 μM/(nA ms) |
| $g_K$ | The conductance of the voltage-gated $K^+$ channels | 26.05897479 mS/cm$^2$ |
| $g_{UNaV}$ | The conductance of the UNaV channels | 6.616093696 mS/cm$^2$ |
| $g_{NaK}$ | The conductance of $Na^+/K^+$ ATPase | 34.43633707 mS/cm$^2$ |
| $g_{Leak}$ | The conductance of the leak channels | 0.05381739 mS/cm$^2$ |
| $g_{Ca}$ | The conductance of the voltage-gated $Ca^{2+}$ channels | 0.010282045 mS/cm$^2$ |
| *x* | Shift of the activation curve | 21.50405795 mV |
| *y* | Shift of the inactivation curve | -9.31006556 mV |

Table S3. Parameters used in Representative parameter sets in the FNAN model.

| Parameter | Description | Value |
|---|---|---|
| C | Membrane Capacitance | 1 μF/cm$^2$ |
| A | Area of Neuron | 0.02 mm$^2$ |
| $V_{Na}$ | Sodium reversal potential | +55 mV |
| $V_K$ | Potassium reverse potential | -100 mV |
| $\tau_{hA}$ | Time-constant of $h_A$ | 15 ms |
| $V_{Ca}$ | Calcium reversal potential | +120 mV |
| Kd | Dissociation constant of KCa channels | 30 μM |
| Ke | Dissociation constant of KNa channels | 32 μM |
| Kf | Parameter in KNa channels | 3 |
| $V_{AMPA}$ | AMPAR reversal potential | 0 mV |
| $V_{NMDA}$ | NMDAR reversal potential | 0 mV |
| $V_{GABA}$ | $GABA_A$R reversal potential | -70 mV |
| $V_{LeNa}$ | Leak $Na^+$ reversal potential | 0 mV |
| $\alpha_{Ca}$ | Coefficient of $Ca^{2+}$ entry | 0.5 μM/(nA ms) |
| $\alpha_{Na}$ | Coefficient of $Na^+$ entry | 1.0 μM/(nA ms) |
| $g_K$ | The conductance of the voltage-gated $K^+$ channels | 19.94160494 mS/cm$^2$ |
| $g_{UNaV}$ | The conductance of the UNaV channels | 0.957341206 mS/cm$^2$ |
| $g_{KNa}$ | The conductance KNa channels | 21.24664 mS/cm$^2$ |
| $g_{Leak}$ | The conductance of the leak channels | 0.052591032 mS/cm$^2$ |
| $g_{Ca}$ | The conductance of the voltage-gated $Ca^{2+}$ channels | 0.068441559 mS/cm$^2$ |

| | | |
|---|---|---|
| $x$ | Shift of the activation curve | 29.80668033 mV |
| $y$ | Shift of the inactivation curve | 10.31760786 mV |
| $g_{Na}$ | The conductance of the voltage-gated $Na^+$ channels | 0.014029333 mS/cm$^2$ |
| $g_{A}$ | The conductance of the fast A-type $K^+$ channels | 0.013216077 mS/cm$^2$ |
| $g_{KS}$ | The conductance of the slowly inactivating $K^+$ channels | 0.170250406 mS/cm$^2$ |
| $g_{KCa}$ | The conductance of the $Ca^{2+}$-dependent $K^+$ channels | 0.47912228 mS/cm$^2$ |
| $g_{NaP}$ | The conductance of the persistent $Na^+$ channels | 0.915345812 mS/cm$^2$ |
| $g_{AR}$ | The conductance of the inwardly rectifying $K^+$ channels | 0.017545974 mS/cm$^2$ |
| $g_{AMPA}$ | The conductance of the AMPA receptors | 9.876574947 μS |
| $g_{NMDA}$ | The conductance of the NMDA receptors | 0.239669033 μS |
| $g_{GABA}$ | The conductance of the $GABA_A$ receptors | 0.161273407 μS |
| $\tau_{Ca}$ | The time constant of $Ca^{2+}$ efflux | 80.08306247 ms |
| $\tau_{Na}$ | The time constant of $Na^+$ efflux | 2272.480552 ms |

Table S4. Parameters used in Representative parameter sets in the Network FNAN model (SWS).

| Parameter | Description | Value |
|---|---|---|
| C | Membrane Capacitance | 1 μF/cm$^2$ |
| A | Area of Neuron | 0.02 mm$^2$ |
| $V_{Na}$ | Sodium reversal potential | +55 mV |
| $V_K$ | Potassium reverse potential | -100 mV |
| $\tau_{hA}$ | Time-constant of $h_A$ | 15 ms |
| $V_{Ca}$ | Calcium reversla potential | +120 mV |
| Kd | Dissociation constant of KCa channels | 30 μM |
| Ke | Dissociation constant of KNa channels | 32 μM |
| Kf | Parameter in KNa channels | 3 |
| $V_{AMPA}$ | AMPAR reversal potential | 0 mV |
| $V_{NMDA}$ | NMDAR reversal potential | 0 mV |
| $V_{GABA}$ | $GABA_AR$ reversal potential | -70mV |
| $V_{LeNa}$ | Leak $Na^+$ reversal potential | 0 mV |
| $\alpha_{Ca}$ | Coefficient of $Ca^{2+}$ entry | 0.5 μM/(nA ms) |
| $\alpha_{Na}$ | Coefficient of $Na^+$ entry | 1.0 μM/(nA ms) |
| $g_K$ | The conductance of the voltage-gated $K^+$ channels | 36.05529836 mS/cm$^2$ |
| $g_{UNaV}$ | The conductance of the UNaV channels | 6.5465182 mS/cm$^2$ |
| $g_{KNa}$ | The conductance KNa channels | 6.841528212 mS/cm$^2$ |
| $g_{Leak}$ | The conductance of the leak channels | 0.015971334 mS/cm$^2$ |

| | | |
|---|---|---|
| $g_{Ca}$ | The conductance of the voltage-gated $Ca^{2+}$ channels | 0.066595773 mS/cm$^2$ |
| $x$ | Shift of the activation curve | 33.33945676 mV |
| $y$ | Shift of the inactivation curve | 12.8051663 mV |
| $g_{Na}$ | The conductance of the voltage-gated $Na^+$ channels | 19.33477692 mS/cm$^2$ |
| $g_{A}$ | The conductance of the fast A-type $K^+$ channels | 0.02762257 mS/cm$^2$ |
| $g_{KS}$ | The conductance of the slowly inactivating $K^+$ channels | 0.089453858 mS/cm$^2$ |
| $g_{KCa}$ | The conductance of the $Ca^{2+}$-dependent $K^+$ channels | 3.056566755 mS/cm$^2$ |
| $g_{NaP}$ | The conductance of the persistent $Na^+$ channels | 1.07002588 mS/cm$^2$ |
| $g_{AR}$ | The conductance of the inwardly rectifying $K^+$ channels | 0.029441142 mS/cm$^2$ |
| $g_{AMPA}$ | The conductance of the AMPA receptors | 0.045167775 μS |
| $g_{NMDA}$ | The conductance of the NMDA receptors | 0.010225012 μS |
| $\tau_{Ca}$ | The time constant of $Ca^{2+}$ efflux | 87.03628253 ms |
| $\tau_{Na}$ | The time constant of $Na^+$ efflux | 1058.482063 ms |

Table S5. Parameters used in Representative parameter sets in the Network FNAN model (AWAKE).

| Parameter | Description | Value |
|---|---|---|
| C | Membrane Capacitance | 1 μF/$cm^2$ |
| A | Area of Neuron | 0.02 $mm^2$ |
| $V_{Na}$ | Sodium reversal potential | +55 mV |
| $V_K$ | Potassium reverse potential | -100 mV |
| $\tau_{hA}$ | Time-constant of $h_A$ | 15 ms |
| $V_{Ca}$ | Calcium reversal potential | +120 mV |
| Kd | Dissociation constant of KCa channels | 30 μM |
| Ke | Dissociation constant of KNa channels | 32 μM |
| Kf | Parameter in KNa channels | 3 |
| $V_{AMPA}$ | AMPAR reversal potential | 0 mV |
| $V_{NMDA}$ | NMDAR reversal potential | 0 mV |
| $V_{GABA}$ | $GABA_AR$ reversal potential | -70 mV |
| $V_{LeNa}$ | Leak $Na^+$ reversal potential | 0 mV |
| $\alpha_{Ca}$ | Coefficient of $Ca^{2+}$ entry | 0.5 μM/(nA ms) |
| $\alpha_{Na}$ | Coefficient of $Na^+$ entry | 1.0 μM/(nA ms) |
| $g_K$ | The conductance of the voltage-gated $K^+$ channels | 36.05529836 mS/$cm^2$ |
| $g_{UNaV}$ | The conductance of the UNaV channels | 6.5465182 mS/$cm^2$ |
| $g_{KNa}$ | The conductance KNa channels | 0.06841528212 mS/$cm^2$ |
| $g_{Leak}$ | The conductance of the leak channels | 0.015971334 mS/$cm^2$ |

| | | |
|---|---|---|
| $g_{Ca}$ | The conductance of the voltage-gated $Ca^{2+}$ channels | 0.066595773 mS/cm$^2$ |
| $x$ | Shift of the activation curve | 33.33945676 mV |
| $y$ | Shift of the inactivation curve | 12.8051663 mV |
| $g_{Na}$ | The conductance of the voltage-gated $Na^+$ channels | 19.33477692 mS/cm$^2$ |
| $g_A$ | The conductance of the fast A-type $K^+$ channels | 0.02762257 mS/cm$^2$ |
| $g_{KS}$ | The conductance of the slowly inactivating $K^+$ channels | 0.089453858 mS/cm$^2$ |
| $g_{KCa}$ | The conductance of the $Ca^{2+}$-dependent $K^+$ channels | 3.056566755 mS/cm$^2$ |
| $g_{NaP}$ | The conductance of the persistent $Na^+$ channels | 1.07002588 mS/cm$^2$ |
| $g_{AR}$ | The conductance of the inwardly rectifying $K^+$ channels | 0.029441142 mS/cm$^2$ |
| $g_{AMPA}$ | The conductance of the AMPA receptors | 0.045167775 μS |
| $g_{NMDA}$ | The conductance of the NMDA receptors | 0.010225012 μS |
| $\tau_{Ca}$ | The time constant of $Ca^{2+}$ efflux | 87.03628253 ms |
| $\tau_{Na}$ | The time constant of $Na^+$ efflux | 1058.482063 ms |
